# *EmbSAM*: Cell boundary localization and Segment Anything Model for fast images of developing embryos

**DOI:** 10.1101/2024.09.07.611795

**Authors:** Cunmin Zhao, Zelin Li, Pei Zhang, Yixuan Chen, Pohao Ye, Ming-Kin Wong, Lu-Yan Chan, Hong Yan, Chao Tang, Guoye Guan, Zhongying Zhao

## Abstract

Cellular shape dynamics are critical for understanding cell fate determination and organogenesis during development. However, fluorescence live-cell images of cell membranes frequently suffer from a low signal-to-noise ratio, especially during long-duration imaging with high spatiotemporal resolutions. This is caused by phototoxicity and photobleaching, which limit laser power and hinder effective time-lapse cell shape reconstruction, particularly in rapidly developing embryos. Here, we devised a new computational framework, *EmbSAM*, that incorporates a deep-learning-based cell boundary localization algorithm and the Segment Anything Model. *EmbSAM* enables accurate and robust three-dimensional (3D) cell membrane segmentation for roundworm *Caenorhabditis elegans* embryos imaged every 10 seconds. The cell shape data prior to gastrulation quantitatively characterizes a series of cell-division-coupled morphodynamics associated with cell position, cell identity, lineage, and fate, and can be accessed locally and online. The framework also exhibits potential in segmenting and quantifying the fluorescence labeling various cell-membraned-attached molecules in both wild-type and RNAi-treated embryos.

## 1. Introduction

Embryogenesis is the developmental process during which a single-celled fertilized egg undergoes cleavage (rapid division) along with the emergence of stereotypical spatial architectures and fate maps of numerous cells, forming a functional larva that can survive independently (Rawlinson 2010, Truman 2019, Xu *et al*. 2020). The rapid division takes place concurrently with cell migration and cell differentiation, where cell shape may change dramatically to facilitate cell fate determination and organogenesis (Pohl *et al*. 2010, Moerman *et al*. 1996). Studying cellular behaviors during rapid division becomes a challenging task. For instance, the duration of cytokinesis (defined between the complete separation of cell nuclei and that of cell membranes) is roughly 2.5 min, during which the cell shape changes from spherical to dumbbell-shaped (Cao *et al*. 2020, Kuang *et al*. 2022). Notably, cytokinesis often exhibits differential behaviors (*e.g*., cell division axis reorientation and time cost) along with specific cell shape dynamics depending on cell positions, cell identities, cell lineages, and cell fates; moreover, these differential behaviors can also exist within a single cell type across successive cell cycles (Sugioka *et al*. 2018, Pimpale *et al*. 2020, Middelkoop *et al*. 2024, Fickentscher *et al*. 2018). The details of cytokinesis and its cell shape dynamics in such a short period are commonly missed in time-lapse 3D imaging due to insufficient spatial and temporal resolutions, especially the temporal one. Moreover, many cellular properties, in particular the ones related to cytokinesis, also change drastically over development, such as the asymmetric partition of cell volume between sister cells and the decrease of cell sphericity within roughly 7 min at late metaphase (Fickentscher *et al*. 2018, Azuma *et al*. 2023). To unravel the biological mechanisms underlying these cellular behaviors, it is essential to acquire fluorescence images of cell membranes through time-lapse 3D live-cell imaging (also called *in toto* imaging) at an exceptional temporal resolution, which are used for cell shape reconstruction. Furthermore, cell shape reconstruction is crucial for quantifying the drastic spatiotemporal dynamics of functional molecules associated with cell membrane and cell division, such as filamentous <u>actin</u> (F-actin) and <u>n</u>on-muscle <u>my</u>osin (NMY) that controls the cell cortex stiffness and fluidity (Sobral *et al*. 2021, Middelkoop *et al*. 2024). However, because of photobleaching and phototoxicity, a tradeoff has to be applied between image quality and laser power or imaging frequency. This is especially true if both the cell identity and cell boundary are simultaneously resolved, with one laser channel for cell nucleus tracing and the other for cell membrane segmentation (Cao *et al*. 2020, Cao *et al*. 2024). Therefore, the development of a cell (membrane) segmentation algorithm has to take into consideration a modest quality of fluorescence image, either for the raw one or processed one.

The transparent roundworm *C. elegans* is one of the most popular models for studying the developmental control over embryogenesis due to its invariant development at the cellular level, including cell lineage, cell migration trajectory, cell division timing and axis orientation, cell volume and fate, and so forth (Fig. S1) (Sulston *et al*. 1983, Ho *et al*. 2015, Guan *et al*. 2019, Guan *et al*. 2021, Packer *et al*. 2019, Ma *et al*. 2021, Wang *et al*. 2022c). Thus, multiple 3D cell segmentation algorithms with time-lapse image stacks of different temporal resolutions have been devised in the past several years: *spheresDT/Mpacts-PiCS* at 3-min intervals (Thiels *et al*. 2021), *CShaper* at ∼1.5-min intervals (Cao *et al*. 2020), and *BCOMS2* at 30-second intervals (Azuma *et al*. 2023).

Apart from the customized frameworks, there are also many techniques developed for universal experimental conditions, in other words, for realizing the cell segmentation with a general computational framework, such as *CellProfiler, RACE*, and *SingleCellDetector* (Soliman 2015, Stegmaier *et al*. 2016, Wang *et al*. 2019). Recently, the Segment Anything Model (*SAM*) based on the <u>Vi</u>sion <u>T</u>ransformer (ViT) architecture further revolutionized the field of computer vision, along with its truncated versions adapted to general 3D biomedical images, *MedLSAM* and *MedSAM* (Dosovitskiy *et al*. 2020, Kirillov *et al*. 2023, Ding *et al*. 2024, Ma *et al*. 2024). The limitations of current *SAM* frameworks include their heavy reliance on manual input for segmentation promoters, such as seeding points or bounding boxes. Although the general *SAM* frameworks with these limitations may not be comparable to the customized ones when executed in the custom system (*e.g*., *spheresDT/Mpacts-PiCS, CShaper*, and *BCOMS2* for the *C. elegans* embryo (Thiels *et al*. 2021, Cao *et al*. 2020, Azuma *et al*. 2023), they still impose a chance that the advantage of different approaches could be integrated so as to increase the overall cell segmentation performance coherently, for example, targeting the fluorescence images with a low <u>s</u>ignal-to-<u>n</u>oise ratio (SNR) just like the ones obtained at a high temporal resolution and with a weak laser power (Guan *et al*. 2019, Guan *et al*. 2021, Kuang *et al*. 2022).

To realize cell-resolved shape reconstruction of developing embryos from fluorescence images with a low SNR, particularly those captured at high temporal resolutions on the order of 10 seconds or less, we devised *EmbSAM*, a computational framework that extends the *SAM* with additional cell boundary localization part containing a denoising module and a watershed module. This framework outputs 3D bounding boxes as a guide to direct the *SAM* to perform precise segmentation of the cell membrane fluorescence in 2D, which can then be assembled in 3D space. Evaluated using three *C. elegans* embryos imaged with a low SNR, *EmbSAM* significantly outperforms *CShaper* and *MedSAM* regarding 3D cell segmentation accuracy and robustness. Furthermore, *EmbSAM* is applied to two more *C. elegans* embryos imaged at 10-second intervals and up to the moment before gastrulation, providing a quantitative measurement of cell shape changes for fundamental cellular behaviors (*e.g*., cell division and cell migration) as well as their dependence on cell positions, cell identities, cell lineages, and cell fates; the resulting data have been reformatted for both local and online analytical platforms previously made available to the public. Developmental landmarks such as post-fertilization pseudocleavage, dorsal-ventral and left-right body axes establishment, and spatial reorganization for gastrulation, are digitized and monitored over the course of time. Our framework also shows potential in segmenting cell membranes with a variety of fluorescently-labeled molecules (*e.g*., the phosphoinositide and non-muscle myosin II), in embryos at different developmental stages and with different genetic backgrounds.

## 2. Methods

### 2.1 Embryo data collection for time-lapse 3D cell segmentation

A total of five *C. elegans* wild-type embryo samples were used for 3D image acquisition and cell segmentation in this paper, with green fluorescence labeling cell nuclei and red fluorescence labeling cell membranes ubiquitously (strain ZZY0535) (Cao *et al*. 2020, Cao *et al*. 2024). The fluorescence labeling on cell nuclei was used for cell lineage tracing and visualization with *StarryNite* (automatic tracing) and *AceTree* (manual correction) respectively (Boyle *et al*. 2006, Bao *et al*. 2006, Murray *et al*. 2006, Santella *et al*. 2010, Santella *et al*. 2010, Katzman *et al*. 2018). Previous fluorescence images were acquired at a relatively low SNR, preventing them from successful 3D cell segmentation by the *CShaper* (one of the most updated algorithms customized for *C. elegans* embryonic images) or *MedSAM* (the most updated Segment Anything Model algorithm generalized for biomedical images) algorithm (Fig. S2) (Cao *et al*. 2020, Ma *et al*. 2024). A total of five embryos were imaged in 3D from no later than the 4-cell stage at two different temporal resolutions as described below.

Three embryo samples were imaged at 1.41-minute intervals for 60 time points, which were imaged in 712×512 pixels on the *xy* plane with a total of 68 focal planes along the *z*-axis (0.09 μm/pixel for the *xy* plane and 0.42 μm/pixel for the *z*-axis, *i.e*., the shooting direction). Two of them were reused from our previous works (Guan *et al*. 2019, Guan *et al*. 2021); these two embryo samples (“Emb1” and “Emb2”) were used for quantitatively evaluating the cell segmentation performance. The last embryo sample (“Emb3”) was first published in this paper, for further independent cell segmentation performance evaluation.

Another two embryo samples were imaged at 10-second intervals. One (“Emb4”) was published in our previous work (Guan *et al*. 2019), which was imaged in 712×512 pixels on the *xy* plane with a total of 47 focal planes along the *z*-axis (0.09 μm/pixel for the *xy* plane and 0.59 μm/pixel for the *z*-axis, *i.e*., the shooting direction) for 260 time points. The other (“Emb5”) was published in our previous work (Kuang *et al*. 2022), which was imaged in 712×512 pixels on the *xy* plane with a total of 47 focal planes along the *z*-axis (0.09 μm/pixel for the *xy* plane and 0.59 μm/pixel for the *z*-axis, *i.e*., the shooting direction) for 300 time points. These two embryo samples exhibit cell cycle lengths strongly proportional to the ones in the embryo samples “Emb1” to “Emb3”, suggesting their fundamental biological process was not affected by the imaging at a significantly higher temporal resolution (almost an order of magnitude) (Fig. S3).

### 2.2. Manual annotation for ground truth

Manual annotation of cell membranes with fluorescence images was used as ground truth, which was necessary for training and evaluating an automatic cell segmentation algorithm. In this work, multiple groups of manually annotated ground truths were used as detailed below.

For training the deep-learning-based denoising module, 16 3D volumetric images manually annotated in five *C. elegans* wild-type embryos (a collection of 2,339 3D cell objects) were adopted from our previous work (Table S1) (Cao *et al*. 2020). They were subsequently sliced in the *x, y*, and *z* directions according to the corresponding recorded pixels, resulting in a total of 4,096, 5,696, and 2,560 2D images, respectively. From these, we randomly selected 10% in each direction to construct the training dataset, which enabled comprehensive learning and adaptation, reflecting the imaging features on fluorescently-labeled cell shapes in different directions.

For evaluating the automatic cell segmentation performance, all the cell shapes of the embryo samples “Emb1” and “Emb2” at 4-, 6-, 7-, 8-, 12-, 14-, 15-, 24-, 26-, 28-, and ≥44-cell stages (a collection of 379 3D cell objects) were meticulously annotated with respect to all the *x, y*, and *z* directions, slice by slice and cell by cell. Besides, a total of 45 2D images (focal planes) in the embryo sample “Emb3” were manually annotated for independent performance evaluation.

### 2.3. Embryo data collection for alternative fluorescently-labeled molecules

For time-lapse imaging, young adult worms were dissected to free embryos. The embryos were mounted on a 3-5% (wt/vol) agarose pad with 0.5% tetramisole and sealed with Vaseline. <u>G</u>reen fluorescent <u>p</u>rotein (GFP) and mCherry were visualized using 488 nm and 561 nm excitation lasers respectively. Channels were imaged sequentially to eliminate bleed-through. Imaging in all channels was captured using 0.1-second exposure time and at 2-to 30-second intervals on an inverted spinning-disk confocal microscope (Olympus SpinSR10) using a Yokogawa CSU W1 scanner system, equipped with a 60×/1.4 NA objective and two Hamamatsu ORCA Flash sCMOS cameras. All movies were acquired under the control of cellSens Dimension software (Olympus), in which multi-*z* sections were merged into a single projected image using ImageJ (Schindelin *et al*. 2012). Images were subsequently arranged using ImageJ with small and global adjustments for contrast and brightness. Embryos with fluorescence labeling the phosphoinositide through pleckstrin homology (PH) domain (“Emb6” to “Emb11”) and <u>n</u>on-muscle <u>my</u>osin <u>II</u> (NMY-2) (“Emb12” to “Emb20”) are involved (Audhya et al. 2005, Matsumura 2005, Lan et al. 2019).

For single-shot imaging, embryos were mounted in M9 buffer with 10 mM sodium azide (Sigma) on glass slides, and observed under the Carl Zesis LSM 980 confocal microscope equipped with a Zeiss 60×/1.40 NA oil immersion objective lens (Carl Zeiss). Lasers 488 nm were used to excite GFP. Single-plane images were taken as 6-10 sections along the *z*-axis at 0.2-μm intervals. Multi-*z* sections were acquired and merged into a single projected image using Zen software (Carl Zeiss). Images were subsequently arranged using Adobe Photoshop with small and global adjustments for contrast and brightness. Embryos with fluorescence labeling <u>f</u>ilamentous <u>actin 5</u> (ACT-5) (“Emb21” to “Emb26”) are involved (Gobel *et al*. 2004, MacQueen *et al*. 2005).

### 2.4. RNA interference

For standard <u>RNA</u> interference (RNAi), about 10 young adults were picked and cultured on RNAi plates (<u>n</u>ematode <u>g</u>rowth <u>m</u>edia (NGM) containing 1 mM iso<u>p</u>ropylthio<u>g</u>alactoside (IPTG) and 100 μg/mL ampicillin) seeded with bacterial clones of target genes, and their first-generation embryos were examined after 72 h. Worms were fed with RNAi bacteria containing the L4440 <u>e</u>mpty <u>v</u>ector plasmid as a control treatment (EV RNAi). All RNAi clones were confirmed by sequencing. For time-lapse images, the young adult worms were put in the M9 buffer, and dissected by two needles to release their embryos. Embryos were then mounted on a 2% agarose pad and imaged with oil immersion objectives.

### 2.5. Proposed cell segmentation framework

The proposed computational framework for cell segmentation, *EmbSAM*, consists of three major parts (Fig. 1):

**Fig. 1.**
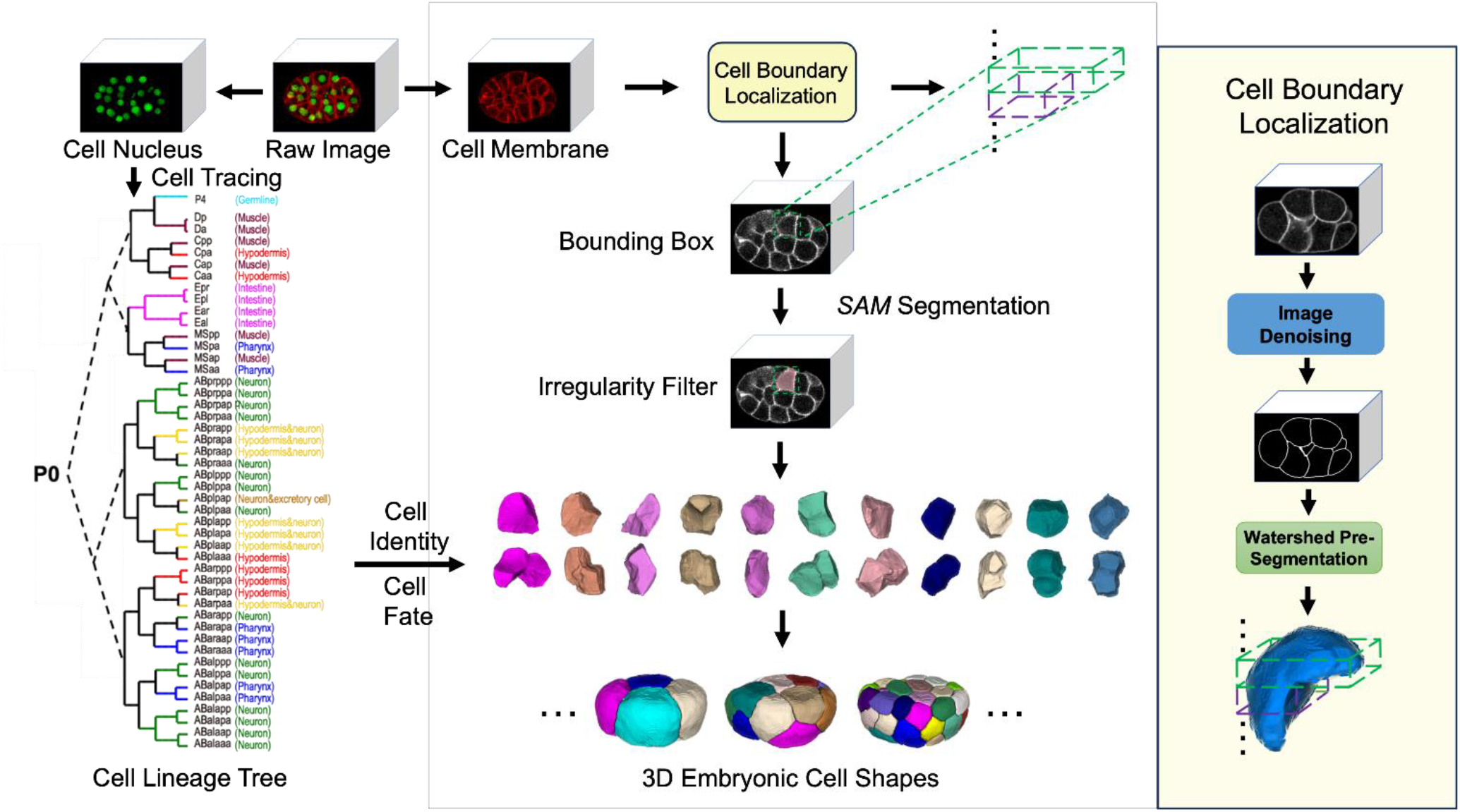
The flowchart of *EmbSAM*.

1. The cell boundary localization part for denoising fluorescence images and generating bounding boxes for each cell region. This is primarily composed of a denoising module (the deep neural network that removes small noisy components to increase the SNR of raw images) and a watershed module (the pre-segmentation for generating bounding boxes based on a watershed algorithm and obtaining approximate boundaries of the target cell to be segmented) (Wang *et al*. 2022a, Cao *et al*. 2019).
2. The Segment Anything Model (*SAM*) part for final automatic cell segmentation (Kirillov *et al*. 2023). This accomplishes automatic cell segmentation based on a series of bounding box promoters. The *SAM* module maximizes the performance of the proposed framework, whose pre-trained model was trained with billions of images containing thousands of imaging conditions. It is one of the reasons that it probably can help deal with low-SNR images. For the target cells in each slice (focal plane), the bounding box produced by the cell boundary location part was used to facilitate the *SAM* segmentation with 3D assembling.
3. The cell tracing part for assigning cell identity to the reconstructed 3D cell regions. The cell nucleus location output by *StarryNite* and *AceTree* is used to match its corresponding 3D cell region.

#### 2.5.1. Image denoising using conditional normalizing flow

The image-denoising network is based on the *LLFlow* (<u>L</u>ow-<u>L</u>ight Image Enhancement with Normalizing <u>Flow</u>) model (Wang *et al*. 2022b), which utilizes a conditional normalizing flow model (Abdelhamed *et al*. 2019, Winkler *et al*. 2019) informed by the Retinex theory (Wei *et al*. 2018). Given a raw image *I*_raw_, the processing procedure includes histogram equalization, color extraction, and noise extraction followed by the *RRDB* (<u>R</u>esidual-in-<u>R</u>esidual <u>D</u>ense <u>B</u>lock) module (Wang *et al*. 2018) for feature extraction, resulting in the illumination invariant color map *G*(*I*_raw_). The trained invertible network *F* of the conditional normalizing flow can construct the transformation process of the probability distribution between the manually annotated ground truth image with low noise (*I*_clean_) and its original cell membrane fluorescence image with high-noise (*I*_raw_) from a latent code *J* that aligns with the standard Gaussian distribution and *G*(*I*_raw_). Here, *F* consists of three layers (*incl*., a squeeze layer and 12 flow steps), with the probability density function *P*(*I* | *I*_ref_) in the condition (reference image) of *I*_ref_ expressed as:

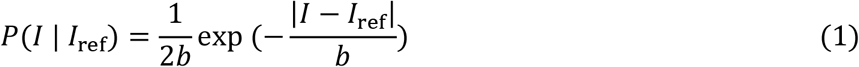

where *b* is a positive constant related to the learning performance. Thus, the probability distribution of *I*_clean_ under the condition of *I*_raw_ can be represented as *P*(*I*_clean_ | *I*_raw_), and the transformation process can be represented as *I*_clean_ = *F*(*J, I*_raw_). From the established conditions, we can derive ∫ *P*(*I*_clean_ | *I*_raw_) ∂*I*_clean_ = ∫ *P*(*J* | *I*_raw_) ∂*J* and *J* = *F*^−1^(*I*_clean_, *I*_raw_). After applying the Jacobian correction to the probability density of *J*, we obtain:

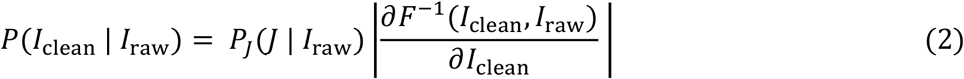

To capture distributional differences between noise and cell features, a negative log-likelihood (NLL) minimization approach is utilized to maximize the probability distribution of *P*(*I*_clean_ | *I*_raw_) to train *F*, getting the loss function:

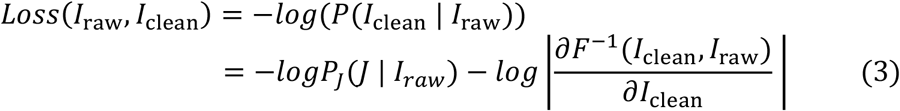

After training, the inference can be implemented onto all raw images beyond the manually annotated ground truths, *Î*_clean_ = *F*(*J, I*_raw_), deriving their low-noise outputs (Fig. 1, Fig. S4).

During denoising, 2D images are stacked along the *z*-axis to construct a 3D image resized to (256, 356, 160) by trilinear interpolation with an even voxel resolution of 0.18 μm/pixel. Then 2D slices are generated by cutting the 3D images along the *x*-, *y*-, and *z*-axis, followed by the denoising process in all three directions. For each pixel in space, the maximum value from its three orthogonal denoised 2D slices is adopted, so that a small-noise 3D image is recombined. This denoising step is essential for taking advantage of the complementary information in three directions regarding the 3D cell membrane fluorescence images. At last, a Gaussian filter is applied for image smoothing, utilizing a Gaussian kernel size of 13 and a standard deviation of 2. The voxel values of the Gaussian-filtered image are constrained to the range (0,1) and then binarized into 0 and 1 using a threshold of 0.5, producing the final 3D image *M*, in which *C* and *E* represent the pixels valued 0 (cell interior and exterior) and 1 (cell boundary) respectively.

#### 2.5.2. Auto-seeding watershed pre-segmentation for generating bounding box promoter

The watershed pre-segmentation algorithm from our previous work (Cao *et al*. 2020) was applied on *M*. Euclidean distance map of *M* is obtained:

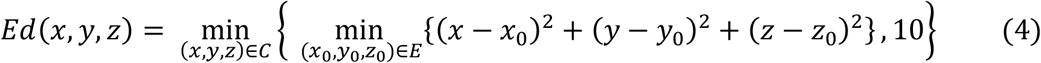

Delaunary triangulation is executed on the local maximum in *Ed* (potential cell centers), where the edge *e*_*ij*_ between vertices *i* and *j* is assigned with a weight by accumulating the *Ed* values along *e*_*ij*_:

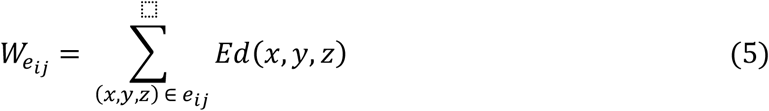

Here, vertices with an edge having a *W* value below a threshold (10) are clustered as the centers inside the same cell (Otsu *et al*. 1979, Cao *et al*. 2019). The vertice clusters (seeding points) and the Euclidean distance map are inputted into the watershed algorithm (Van Der Walt *et al*. 2014), marking both foreground (cell interior) and background (cell exterior) regions. Upon completion of these steps, the binary image with only the embryo interior and exterior is converted into the one with distinct 3D cell regions (Fig. 1, Fig. S4). For each *z* slice in a pre-segmented 3D cell region, the rectangle enclosing its 2D cell region (represented as [*x*_min_: *x*_max_, *y*_min_: *y*_max_, *z*_*i*_]) forms a series of 2D bounding box promoters for the following *SAM* module.

#### 2.5.3. Segment Anything Model

For each time point, all the 2D bounding boxes surrounding a specific cell outputted by the watershed module were fed into the pre-trained zero-shot Segment Anything Model (base vision model named vit_b) (Kirillov *et al*. 2023). The segmentation output of *SAM* is a 3D cell region comprising both background (0, cell exterior) and foreground (1, cell interior).

When performing cell segmentation, the *SAM* module might generate multiple potential 2D regions within the same area, including noises, the target cell, or its neighboring cells. Here, we selected the largest area as the target cell region. Additionally, to exclude unreasonable irregular regions, such as dispersed coralloid or starlike shapes, we calculated the irregularity of each 2D cell region, 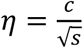, where *c* and *s* represent its circumference and surface area respectively. For calculating a 2D cell region’s perimeter, we used the Douglas-Peucker algorithm for contour approximation, with the approximation coefficient set as 0.01 (Douglas *et al*. 1973). By analyzing the irregularity of 265,704 manually annotated 2D cell regions in our previous works (Cao *et al*. 2020, Guan *et al*. 2023), a threshold value of 9.02 was obtained for establishing a reasonable range for 2D cell irregularity at the 99% confidence (Fig. S5). After filtering by the cell irregularity threshold, all the remaining reliable 2D regions of a target cell are assembled into a 3D region, so that cell-resolved shape reconstruction over embryonic development is achieved.

### 2.6. Cell segmentation performance evaluation

To systematically assess the similarity between the cell segmentation results and their ground truth annotations, we adopted two widely recognized metrics for 3D object comparison (Huttenlocher *et al*. 1993, Taha *et al*. 2015) :

- Dice score: The ratio of the overlapping volume to the total volume of two 3D objects.
- Hausdorff distance: The maximum distance calculated from every voxel in one 3D object to its nearest voxel in another 3D object.

In theory, a larger Dice score and a smaller Hausdorff distance indicate a higher consistency between the cell segmentation results and their ground truth annotations.

### 2.7. 3D cell shape descriptor

The characteristics of cell shapes enclosed by their segmented cell membranes can be quantitatively described by a series of 3D shape descriptors with explicit geometric significance. Here, we adopted three 3D cell shape descriptors from our previous work and calculated a new one as follows (Guan *et al*. 2024).

- Taking a perfect sphere with the same volume as the cell, “*general sphericity*” is defined as the ratio of its surface area to that of the cell, in other words, it describes the similarity of the cell to a perfect sphere (Wadell 1932, Cruz-Matías *et al*. 2019):

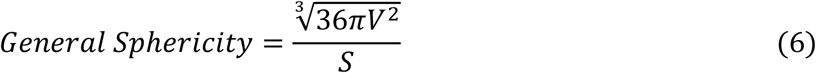

where *V* and *S* are the volume and surface area of the cell respectively.
- While “*general sphericity*” assesses the gross shape of a cell, “*Hayakawa roundness*” specifically assesses the sharpness of edges and corners, as well as the presence of the convexities and concavities on the cell surface (Cruz-Matías *et al*. 2019, Hayakawa *et al*. 2005):

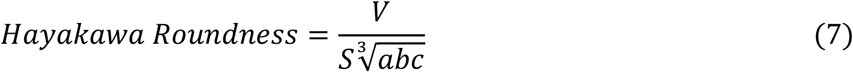

where *a, b* and *c* are the length of the long, intermediate, and short axes of the <u>o</u>riented <u>b</u>ounding <u>b</u>ox (OBB) of the cell, respectively, estimated by principal component analysis (Cruz-Matías *et al*. 2019, Zhao *et al*. 2017).
- Derived from a 2D definition, “*spreading index*” reflects the degree to which the convex hull of a cell resembles a perfect sphere, *i.e*., the spreading of the cell shape: (Yu *et al*. 2013, Lobo *et al*. 2016):

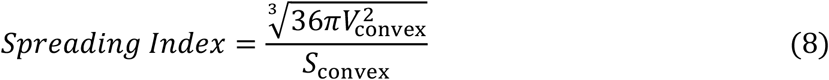

where *V*_convex_ and *S*_convex_ are the volume and surface area of the convex hull enclosing the cell respectively.
- During cytokinesis, the mother cell nucleus divides into two daughter cell nuclei with locations (*x*_nuc1_, *y*_nuc1_, *z*_nuc1_) and (*x*_nuc2_, *y*_nuc2_, *z*_nuc2_); almost at the same time, the cell membrane elongates with the equatorial plate ingressing as a contractile ring, whose diameter keeps shrinking (Taneja *et al*. 2020). The equatorial plate with a contractile ring is presumed as perpendicular to the line between the two daughter cell nuclei. Hence, the plane equation is described by (*Ax* + *By* + *Cz* + *D* = 0), where (*A, B, C*) = (*x*_nuc2_ − *x*_nuc1_, *y*_nuc2_ − *y*_nuc1_, *z*_nuc2_ − *z*_nuc1_) is its normal vector defined by the locations of the two daughter cell nuclei and *D* is determined as follows. All pixels with a distance to the plane smaller than 0.5 pixels form an approximate cylinder with a height of one pixel, by which the diameter of the contractile ring can be derived from its surface area:

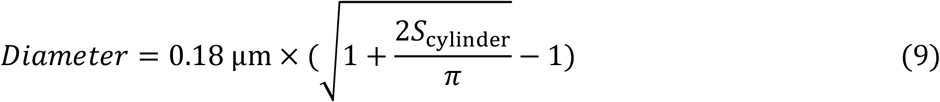

where *S*_cylinder_ is the surface area of the convex hull enclosing the cylinder and *D* is determined by searching the smallest *S*_cylinder_.

## 3. Results

### 3.1. Performance of EmbSAM in cell segmentation on low-SNR images

With *C. elegans* wild-type embryo samples “Emb1” and “Emb2” at their 4-, 6-, 7-, 8-, 12-, 14-, 15-, 24-, 26-, 28-, and ≥44-cell stages, we compared *EmbSAM* to *CShaper* (one of the most updated algorithms customized for *C. elegans* embryonic images) and *MedSAM* (the most updated Segment Anything Model algorithm generalized for biomedical images) (Cao *et al*. 2020, Cao *et al*. 2024, Ma *et al*. 2024). Considering that *MedSAM* was designed for segmenting 2D images and demands bounding box promoters to guide the reconstruction of 3D objects, the cell nucleus location (*x*_nuc_, *y*_nuc_, *z*_nuc_) and the conserved *C. elegans* embryonic cell volume (*V*) documented before were utilized for constructing the required 3D bounding box promoters, a cuboid with boundaries 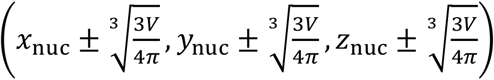 (Cao *et al*. 2020, Cao *et al*. 2024); therefore, the segmented 2D regions from *MedSAM* can be transformed into 3D regions. Intuitively, the *EmbSAM* segmentation outputs smooth and compacted cell shapes in both 2D and 3D for the whole embryo at developmental stages with a few to dozens of cells (Fig. 2), while the segmented cell shapes from *CShaper* and *MedSAM* are coarse or uncompacted (Fig. S2).

**Fig. 2.**
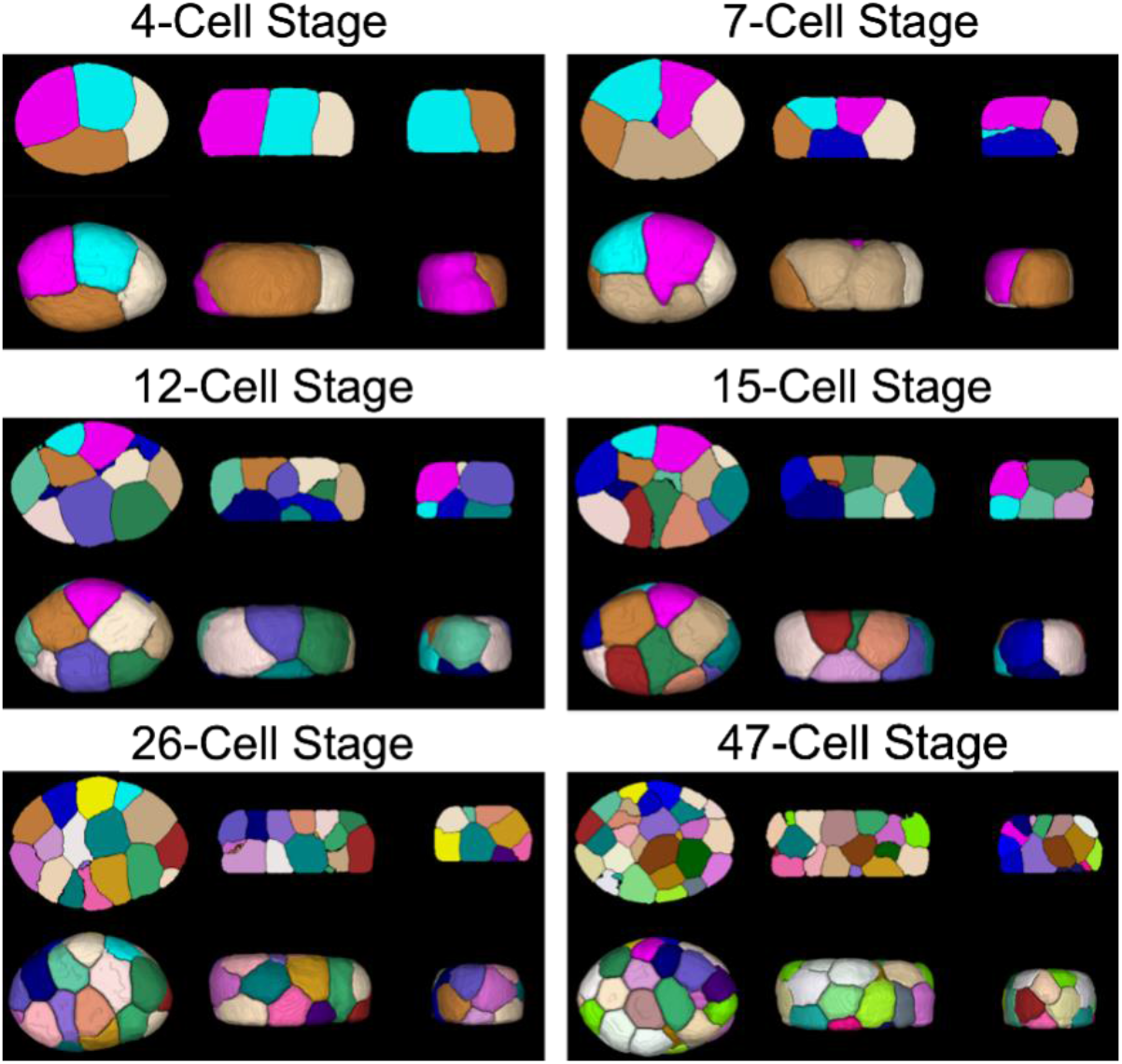
The segmentation results (exemplified by the embryo sample “Emb1”) of *EmbSAM* viewed in different perspectives at different developmental stages. For each developmental stage, the middle 2D crosssection and overall 3D illustration are viewed in the top and bottom rows respectively, while the views along the *z*-, *y*-, and *x*-axis are shown from left to right accordingly.

While quantitative evaluation shows that *CShaper* and *MedSAM* (Cao *et al*. 2020, Cao *et al*. 2024, Ma *et al*. 2024) have similar performance at specific developmental stages like 12- and ≥44-cell stages, *EmbSAM* achieves both the largest Dice score and smallest Hausdorff distance on average across all developmental stages, supporting its outperformance in segmentation accuracy; in addition, *EmbSAM* always achieves the smallest standard deviation in both metrics, supporting its outperformance in segmentation robustness (Fig. 3A). Compared to the coarse or uncompacted cell shapes outputted by *CShaper* and *MedSAM*, the ones by *EmbSAM* resemble those of the ground truth (Fig. 3B). The segmentation accuracy and robustness were further demonstrated using the additional embryo sample “Emb3” first published in this work (Fig. S6).

**Fig. 3.**
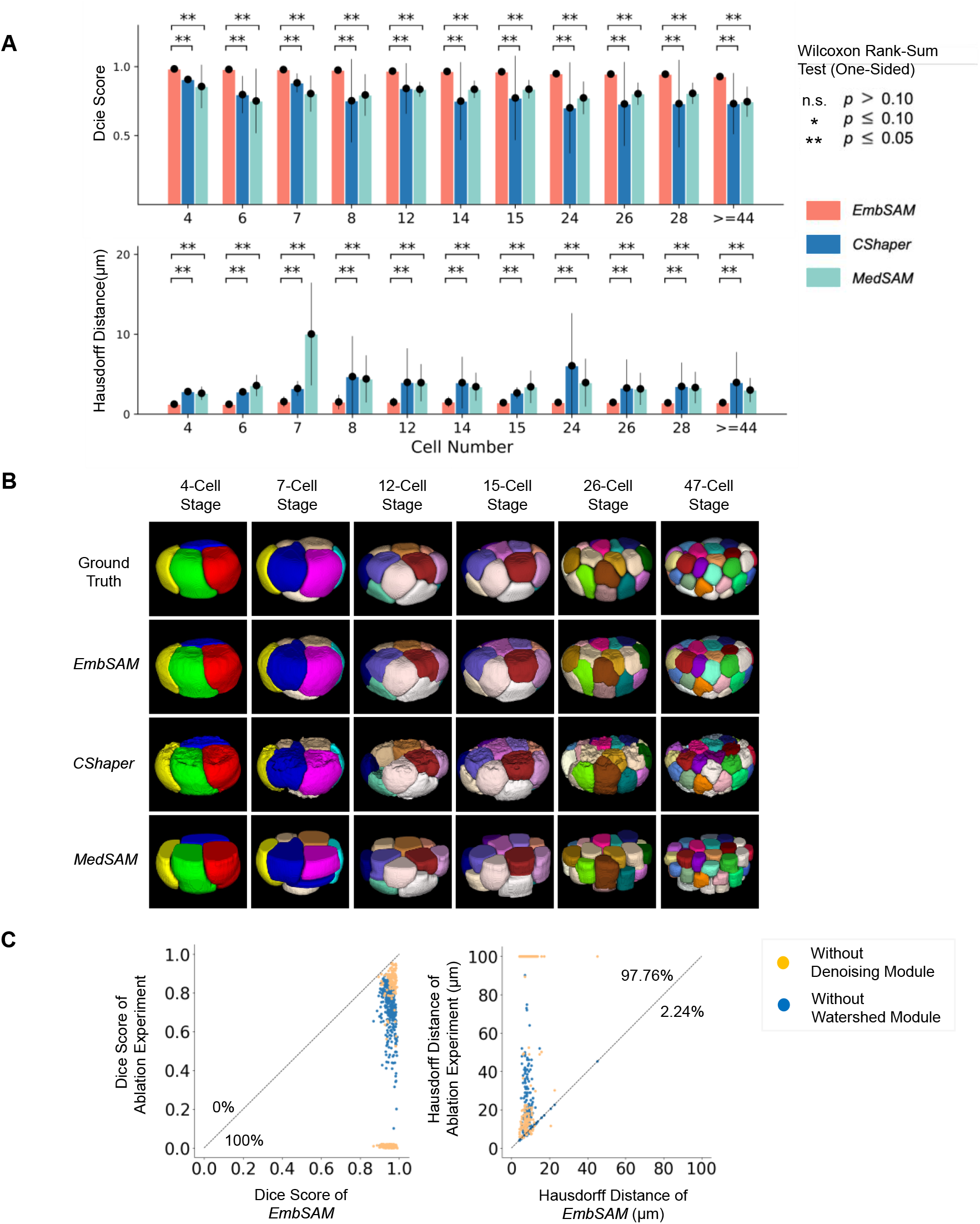
The segmentation performance of *EmbSAM*. (**A**) Outperformance of *EmbSAM* compared to *CShaper* and *MedSAM*, revealed by Dice score (top) and Hausdorff distance (bottom). Statistical significance (one-sided Wilcoxon rank-sum test): n.s. (not significant), *p* > 0.10; *, *p* < 0.10; **, *p* < 0.05; ***, *p* < 0.01. (**B**) Segmentation results (exemplified by the embryo sample “Emb1”) of ground truth (1^st^ row), *EmbSAM* (2^nd^ row), *CShaper* (3^rd^ row), and *MedSAM* (4^th^ row) at different developmental stages. Shown are side views with the anterior of the embryo to the right. (**C**) Poorer segmentation performance of *EmbSAM* when the denoising module and watershed module are ablated respectively, revealed by Dice score (left) and Hausdorff distance (right). The percentage of 3D cell regions with increasing or decreasing values is marked near the diagonal line.

### 3.2. Substantial contribution of denoising module and watershed module

To validate the effectiveness and necessity of the denoising module and watershed module before *SAM* in the *EmbSAM* framework, we implemented an ablation experiment for each of them. For the ablation experiment with the watershed module removed, the cuboid with boundaries 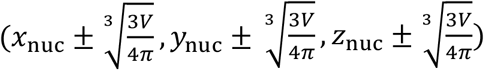 surrounding the cell nucleus location is used as the 3D bounding box promoters for *SAM* segmentation. The ablation experiments reveal a global decline in segmentation performance following the removal of each module (Fig. 3C). While 97.76% of the 3D cell regions acquired poorer Hausdorff distance after the denoising module removal, all of them acquired poorer Dice score. This evidence strongly supports the pivotal role of these two modules in recognizing individual cells in noisy images and promoting the accuracy and robustness of *SAM* segmentation. Such severe segmentation defects in ablation experiments were mostly seen at the top and bottom of the cells (Fig. S7), where the fluorescence signal intensity is relatively lower due to the single-layer cell membrane (compared to the double-layer ones formed by two contacting cells inside the embryo) and the loss of laser intensity through the *z*-axis (parallel to the shooting direction and perpendicular to the focal plane) in the embryo (Guan *et al*. 2022).

### 3.3. Monitoring 3D morphodynamics of cell biology event at 10-second intervals

Embryonic cell divisions proceed with drastic cell shape dynamics as fast as seconds to minutes, such as cytokinesis, which is coupled with rapid cell motion, asymmetric cell volume segregation, and fate differentiation (Fickentscher *et al*. 2017, Guan *et al*. 2021, Pimpale *et al*. 2020, Sugioka *et al*. 2018, Fickentscher *et al*. 2018). Since the *EmbSAM* framework can effectively segment the cell membrane fluorescence images with a low SNR in the embryo samples “Emb1” to “Emb3” (Fig. 2, Fig. 3, Fig. S6, Fig. S7), we further applied it to another two *C. elegans* wild-type embryo samples, “Emb4” and “Emb5”, that were imaged at a high temporal resolution but with a weak laser intensity (Kuang *et al*. 2022, Guan *et al*. 2019). Fascinatingly, the overall 3D cell shapes inside both embryos were successfully reconstructed up to the moment before gastrulation (*i.e*., 26-cell stage) at 10-second intervals (Movie S1, Movie S2), allowing a detailed study of specific cellular behaviors and developmental landmarks with traced cell identities, lineages, and fates, as shown below (Nance *et al*. 2002, Nance *et al*. 2005, Girard *et al*. 2007). The digital embryonic cell shape data has been reformatted for convenient access, visualization, and analysis through public platforms, including both the local software *ITK-SNAP-CVE* and the online website https://bcc.ee.cityu.edu.hk/cmos/embsam (Movie S3, Movie S4) (Guan et al. 2023).

#### 3.3.1. Cell shape dynamics related to cell division

The cell division orientation and cell cycle length have been known to be regulated by various biomechanical and biochemical processes (Sugioka *et al*. 2018, Fickentscher *et al*. 2018). Our segmented 3D cell shape data can illustrate the cell divisions in multiple lineages and generations at 10-second intervals, exemplified by the AB cells (the 1^st^ somatic founder cell derived from the 1^st^ cell division post fertilization) (Fig. S8), EMS cell (the 2^nd^ somatic founder cell derived from the 3^rd^ cell division post fertilization) (Fig. 4A), MS and E cells (anterior and posterior daughter cell of EMS) (Fig. S9), and C and P3 cells (the 3^rd^ somatic founder cell and remaining germline stem cell derived from the 7^th^ cell division post fertilization) (Fig. S10). Taking the EMS cell division for a case study, the Wnt signaling from its neighbor cell, *i.e*., the P2 cell, controls its axis orientation and differentiation of the two daughter cells, *i.e*., the MS cell for mesoderm and the E cell for endoderm (Fig. 4A, Movie S5) (Thorpe *et al*. 1997, Rocheleau *et al*. 1997). Both the separation and motion of cell nuclei and cell membrane can be vividly visualized and quantitatively characterized, at the temporal resolution three times of the previous one (Fig. 4A, Fig. S11) (Azuma *et al*. 2023). At the end of EMS cell division, in other words, the anaphase (starting with a symbol of cell nuclei separation), an increase of cell surface area is detected, in consistency with previous cell biology knowledge on cell division (Sanger *et al*. 1984, Tanaka *et al*. 2020); the majority of increased cell surface area is added to EMS’s contact with the AB cells located in the left of the embryo, ABal and ABpl, indicating the participation of EMS cell division in left-right symmetry breaking during *C. elegans* embryogenesis (Fig. 4BC). Meanwhile, the cell sphericity keeps declining as reported before (Azuma *et al*. 2023), along with the other three independent cell shape descriptors (*i.e*., Hayakawa roundness, spreading index, and diameter) obeying the same trend (Fig. 4D) (Guan *et al*. 2024).

**Fig. 4.**
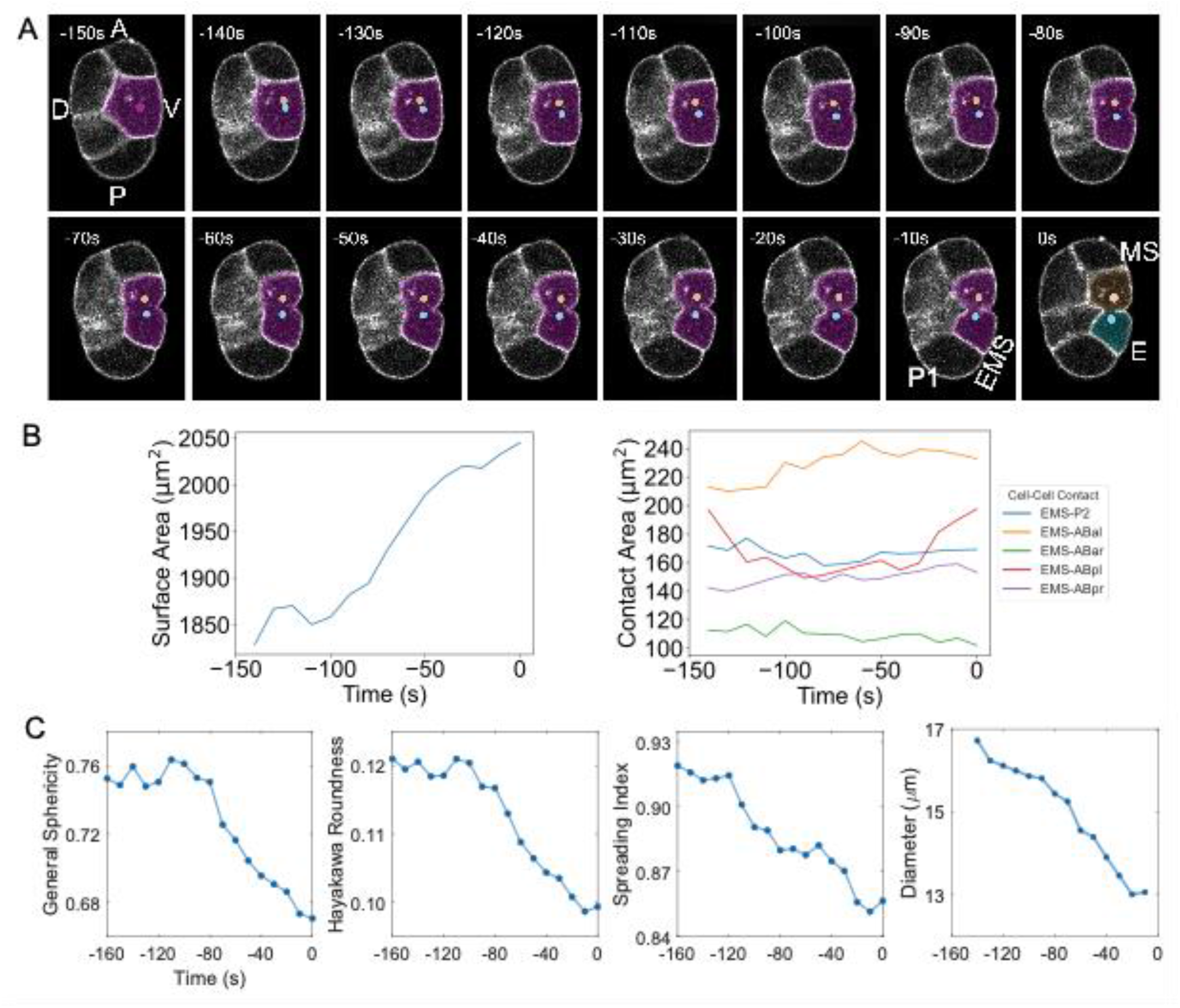
The segmentation results (exemplified by the embryo sample “Emb5”) of *EmbSAM* for the EMS cell division at 10-second intervals. (**A**) 2D segmentation results viewed in the shooting direction and highlighted by the dotted cell nuclei and masked cell membranes. Shown are lateral views with the anterior of the embryo to the top. (**B**) Curves of cell surface area and cell-cell contact areas of the EMS cell. (**C**) Monotonically declining curves of cell shape descriptors for the EMS cell over time. For (**A**)(**B**)(**C**), the absolute developmental time is shown, with the moment of complete cell membrane separation as time zero.

The cell deformation with declining general sphericity, Hayakawa roundness, spreading index, and diameter during cell division is actually proceeding with the cell shape changed from spherical to dumbbell-shaped (Fig. 4AC). Exemplified by the ABpl, ABpr, E cells and their neighbor cells (MS, C, ABplp correspondingly), when a cell initiates its division program, it firstly turns spherical with its interface protruding toward its neighbor cell, and subsequently turns to dumbbell-shaped squeezing its neighbor cell severely into a flat shape (Fig. 5). This implies a strong intracellular and intercellular mechanical force generated by the dividing cell. Such an intensive passive force and deformation exerted by a dividing cell on its neighbor cell appear to be a common phenomenon, further supported not only by the previously reported EMS and P2 cells (squeezed by ABa and ABp cell divisions) but also by other cells – MS (squeezed by ABpl cell division), and E (squeezed by ABpl cell division), C (squeezed by ABpr cell division),(Fig. S12, Fig. S13, Movie S6) (Guan *et al*. 2023). Previous fluorescence imaging on the spatiotemporal dynamics of F-actin demonstrates its accumulation on the cell membrane near cell division and in the cytosol beyond cell division, which makes the cells get rounder and stiffer when it is around cell division (further validated by atomic force microscopy in parallel) (Fujii *et al*. 2021). Our data can clearly illustrate such cell cycle-dependent cell shape dynamics at 10-second intervals, represented by the cells mentioned above when they turn spherical and dumbbell-shaped successively, squeezing their neighbor cells (relatively soft) to adapt to it (relatively hard) through severe deformation.

**Fig. 5.**
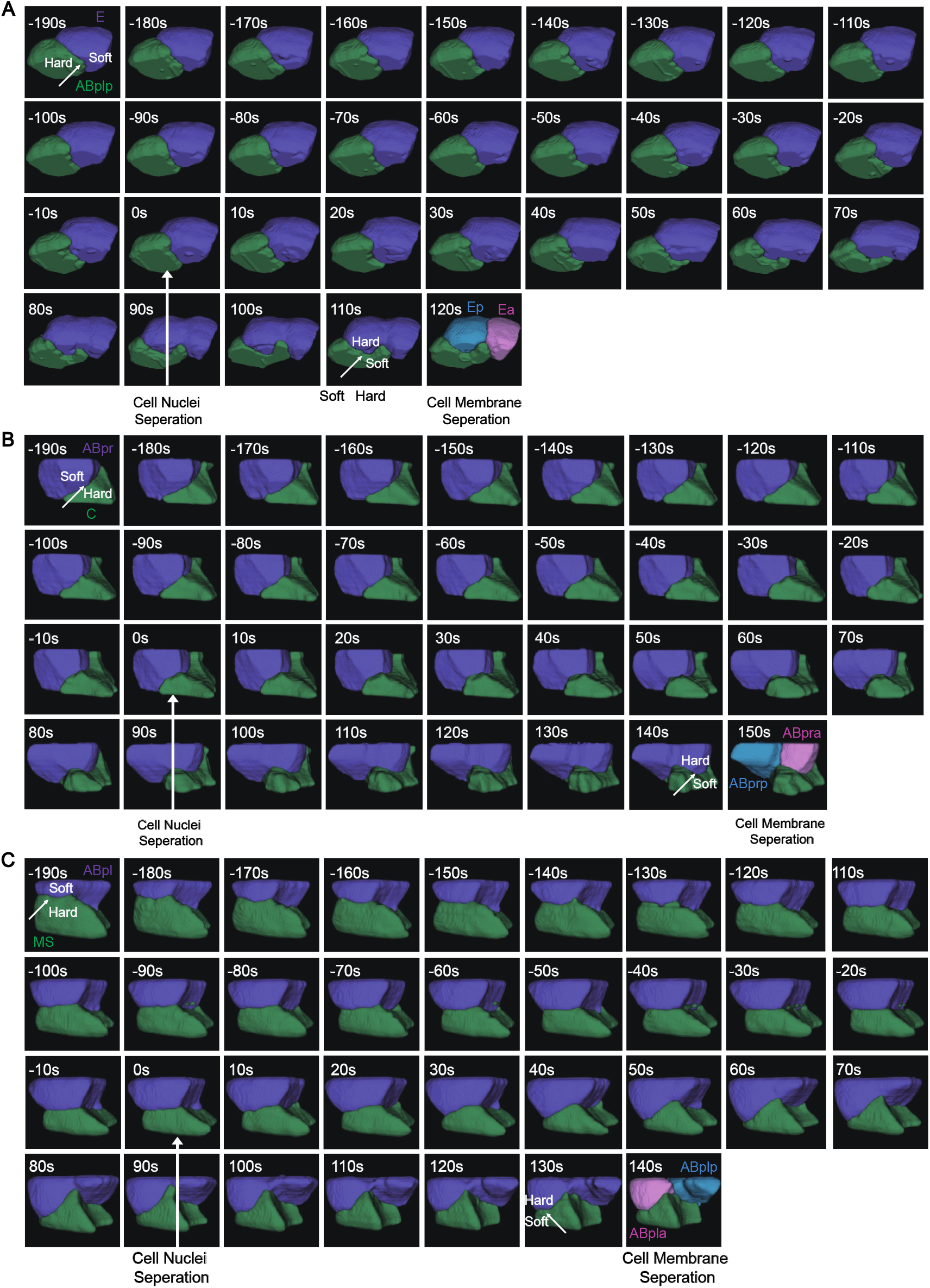
The segmentation results of *EmbSAM* for the drastic shape dynamics of dividing cells (purple), their neighbors (green), and newborn daughters (pink and blue) at 10-second intervals. (**A**) E cell division. (**B**) ABpr cell division. (**C**) ABpl cell division. For (**A**)(**B**)(**C**), the absolute developmental time is shown, with the moment of complete cell nuclei separation as time zero.

#### 3.3.2. Cell shape dynamics related to body axis establishment

The anaphase and telophase of cell division, defined as starting from cell nuclei separation and ending in cell membrane separation, is as fast as 2.5 min measured before (Cao *et al*. 2020, Kuang *et al*. 2022). Such a short-term biological process is critical to establishing the body axes that determine the <u>d</u>orsal (D), <u>v</u>entral (V), left (L), and right (R) of an embryo (also called symmetry breaking), while the <u>a</u>nterior-<u>p</u>osterior (A-P) axis is determined by sperm entry and cell polarization that makes the first cell division asymmetric (Goldstein *et al*. 1996, Motegi *et al*. 2011). While the first two cells AB and P are located in the anterior and posterior of the embryo respectively, the second cell division taking place in AB is initiated with an axis perpendicular to the A-P axis first and then reoriented to it, making its posterior daughter ABp determining the dorsal of the embryo (Fig. 6A, Movie S6).

**Fig. 6.**
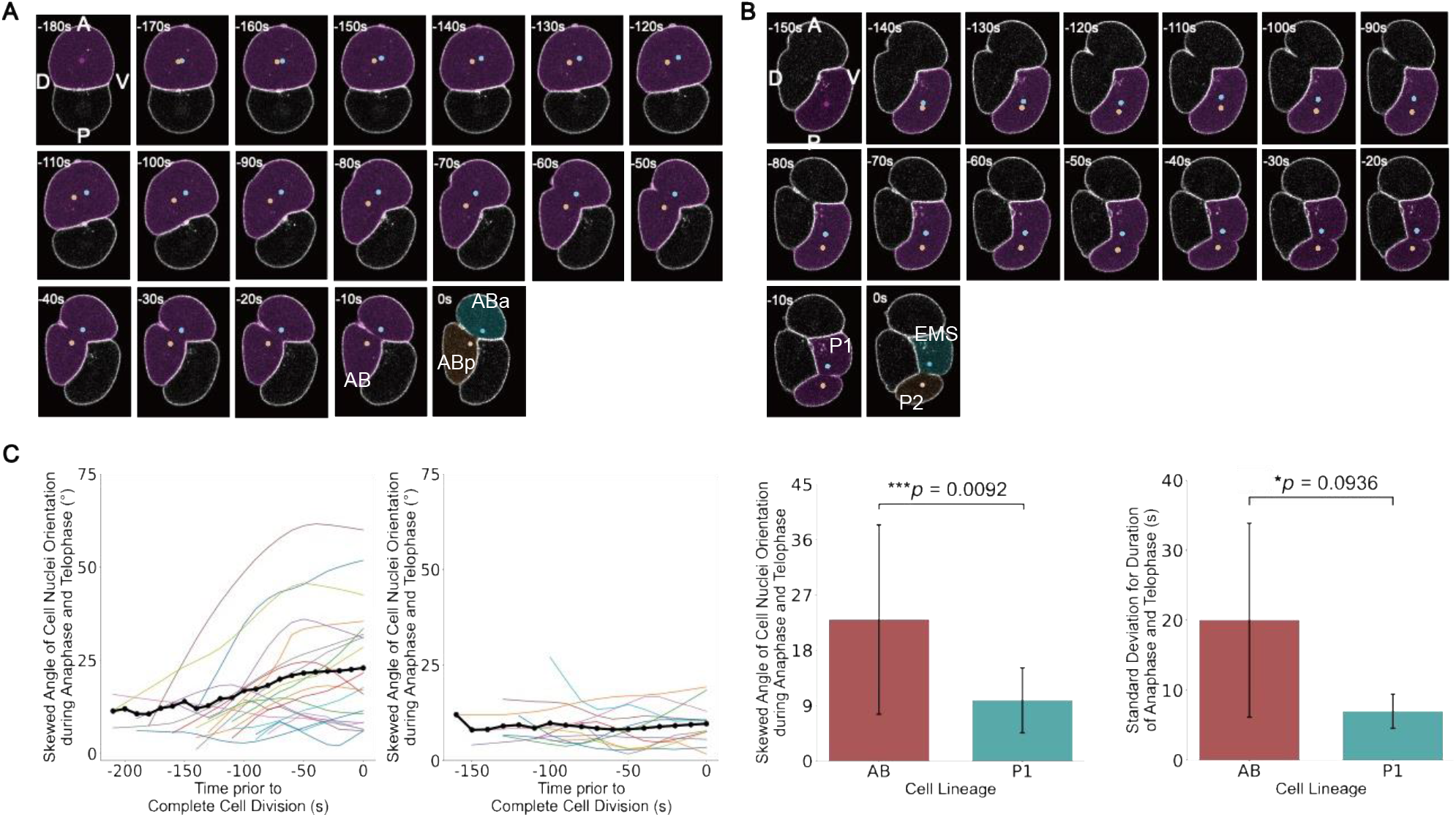
The segmentation results (exemplified by the embryo sample “Emb5”) of *EmbSAM* for the cell division responsible for dorsal-ventral axis establishment at 10-second intervals. (**A**) 2D segmentation results of the AB cell division with the large skewed angle of cell nuclei orientation, viewed in the shooting direction and highlighted by the dotted cell nuclei and masked cell membranes. Shown are lateral views with the anterior of the embryo to the top. (**B**) 2D segmentation results of the P1 cell division with the small skewed angle of cell nuclei orientation, viewed in the shooting direction and highlighted by the dotted cell nuclei and masked cell membranes. Shown are lateral views with the anterior of the embryo to the top. For (**A**)(**B**), the absolute developmental time is shown, with the moment of complete cell membrane separation as time zero. (**C**) Curves of the skewed angle of cell nuclei orientation of all recorded cells in the AB lineage (left) and P1 lineage (right). (**D**) Comparison between all recorded cells in the AB lineage and P1 lineage, regarding the skewed angle of cell nuclei orientation during anaphase and telophase (left), and the standard deviation for duration of anaphase and telophase (right). Statistical significance (independent *t*-test): n.s. (not significant), *p* > 0.10; *, *p* < 0.10; **, *p* < 0.05; ***, *p* < 0.01.

As previous experimental observation indicated that the cell divisions in the AB lineage have a regulated axis reorientation during cytokinesis while the ones in the P1 lineage don’t (Fig. 6B) (Pimpale *et al*. 2020), we measure the skewed angle of cell nuclei orientation during anaphase and telophase for all cells recorded. Shown by the average curve at the temporal resolution of 10 seconds, the AB cells exhibit a stably increasing skewed angle deviated from its initial direction but the P1 cells exhibit a stably unchanged value, faithfully supporting previous conclusion (Fig. 6C). Although the AB cells carry substantially larger skewed angle than P2 cells on average, the standard deviation is also larger (Fig. 6D), meaning that the underlying regulated mechanism may have a variation among the AB cells; this variation might be an intrinsic noise or programmed differential behaviors, worth investigation in the future using our data coupled with cell identities, cell lineages, and cell fates.

Following the diamond-shaped 4-cell stage with both anterior-posterior and dorsal-ventral symmetry breaking, the fourth and fifth cell divisions simultaneously taking place in ABa and ABp (anterior and posterior daughter of AB) are initiated with an axis roughly perpendicular to the plane constituted by the A-P and D-V axes; regulated by a contact-induced myosin flow demonstrated before (Sugioka *et al*. 2018), the axis is slightly skewed with the left daughter cells nearer to the anterior (Fig. 7A, Movies S7). Subsequently, the ABpl cell (left daughter of ABp) undergoes long-range migration toward the dorsal of the embryo with its migration-coupled spreading shape occurring in the middle of its lifespan (Pohl *et al*. 2010, Guan *et al*. 2024). The migration of ABpl, which has been identified with the longest distance among all cells before the 24-cell stage and has nearly the most irregular shape among all cells before the 350-cell stage (Guan *et al*. 2019, Cao *et al*. 2020), was proposed to be driven by cell adhesion regulation (Dutta *et al*. 2019, Kuang *et al*. 2022) and enhances the left-right symmetry breaking substantially. Interestingly, although the duration of anaphase and telophase is revealed to be positively associated with cell volume (Fig. 7C), the one of ABpl is significantly shorter than those of its sister and cousins with similar size (relative difference smaller than 8%), which is probably caused by its varied cytoskeleton state (Fig. 7D). In the future, one might analyze the spatial distribution of functional subcellular structures (*i.e*., lamellipodia, protrusion, and filopodia) over time to understand the underlying mechanism for cell migration, how the migrating cell interacts with its neighbors, and how are different cellular properties (*e.g*., cell migration, cell shape, cell cycle, cell identity, cell lineage, and cell fate) affect each other.

**Fig. 7.**
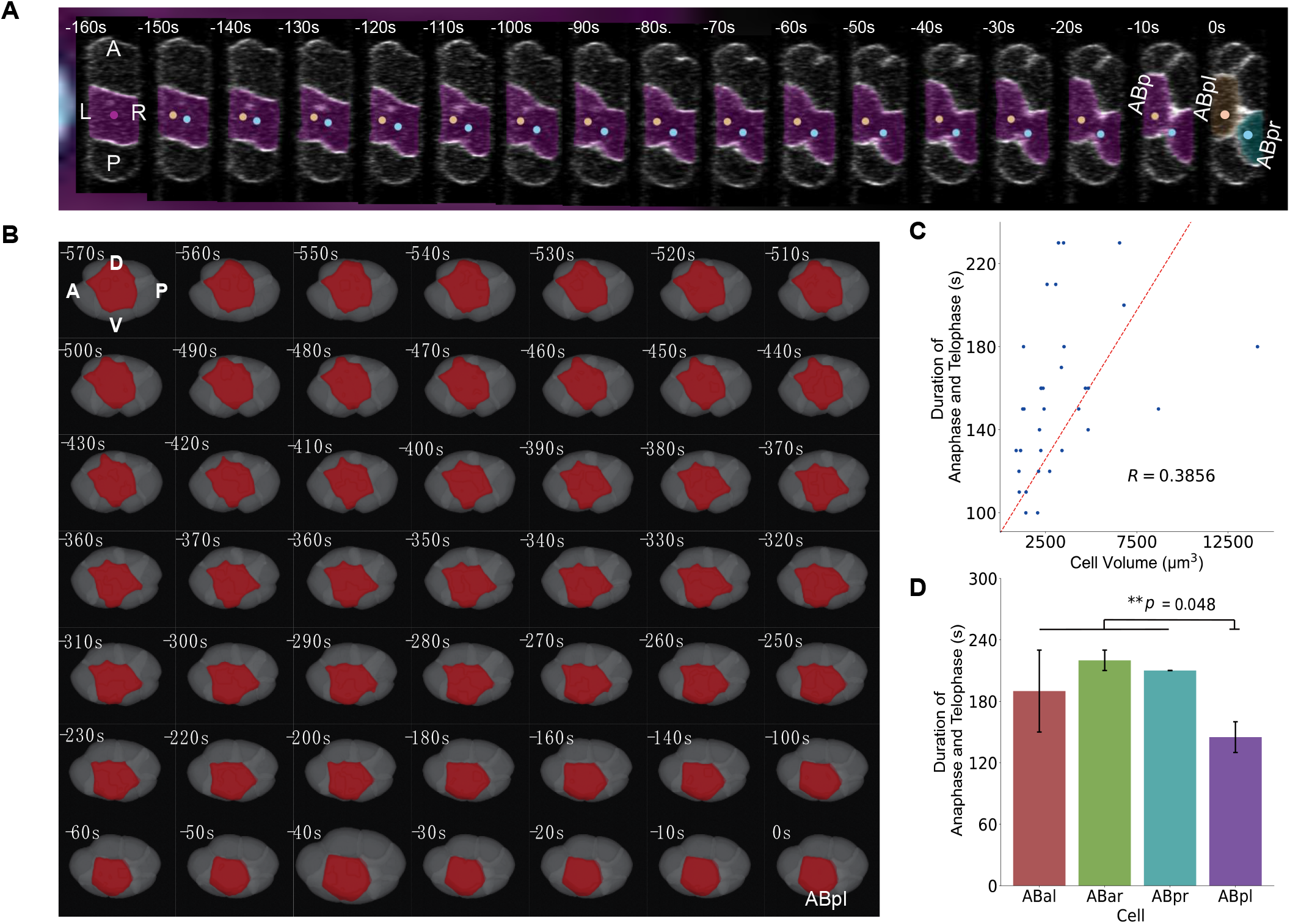
The segmentation results (exemplified by the embryo sample “Emb5”) of *EmbSAM* for the cell division responsible for left-right axis establishment at 10-second intervals. (**A**) 2D segmentation results of the ABp cell division, observed from the dorsal view and highlighted by the dotted cell nuclei and masked cell membranes. Shown are lateral views with the anterior of embryo to the top. The absolute developmental time is shown, with the moment of complete cell membrane separation as time zero. (**B**) 3D segmentation results (exemplified by the embryo sample “Emb4”) of *EmbSAM* for the ABpl cell migration at 10-second intervals. Shown are lateral views with the anterior of the embryo to the left. ABpl is colored in red and other cells are in gray. The absolute developmental time is shown, with the last moment before ABpl cell nuclei separation as time zero. (**C**) Positive correlation between cell volume and the duration of anaphase and telophase with a modest linear correlation coefficient. (**D**) Significance comparison between ABpl cell and its sister and cousins (*i.e*., the ABal, ABar, and ABpr cells), regarding the duration of anaphase and telophase. Statistical significance (independent *t*-test): n.s. (not significant), *p* > 0.10; *, *p* < 0.10; **, *p* < 0.05; ***, *p* < 0.01.

#### 3.3.3 Cell shape dynamics related to spatial reorganization for gastrulation

In addition to morphogenetic events driven by one or a few cell divisions, the ones proceeding over multiple rounds of cell divisions can also be characterized quantitatively at exceptional spatiotemporal resolutions. Previous experimental studies have reported that *C. elegans* early embryonic cells undergo spatial reorganization for gastrulation through apical-basal polarization, where all cells remain attached to the eggshell and form a cavity (blastocoel) to facilitate the upcoming cell ingression, *i.e*., gastrulation (Nance *et al*. 2002, Ajduk *et al*. 2016). In our data with a temporal resolution of 10 seconds, the cells keep acquiring enlarged lateral contact area and slimmer shape to get aligned on the inner surface of the eggshell regularly (Fig. S14A). This is supplemented by a strong negative correlation observed between the relative outer surface area (contacting the eggshell) of a cell and developmental time (Fig. S14B).

### 3.4. Alternative fluorescently-labeled molecules for cell membrane segmentation in embryos with different genetic backgrounds

Although the homogeneously-distributed molecule (*e.g*., phosphoinositide) has been widely used for cell membrane fluorescence labeling, we wondered if it can be replaced by the others with heterogeneous distribution and physiological function of interest. For simplicity, we focused on the 1-to 4-cell stage when all cells are located within the APDV plane and examined if the projected image can be segmented by *SAM*, using phosphoinositide and NMY-2 (<u>n</u>on-muscle <u>my</u>osin <u>II</u>) as comparable fluorescently-labeled targets (Table S2) (Audhya *et al*. 2005, Lan *et al*. 2019). Remarkably, segmentation performs well with clear cell membrane dynamics digitalized. The pseudoceavage (occurring soon after sperm entry with the cell membrane attempting to divide but recovering halfway) and first cell division are reconstructed virtually by both embryos with phosphoinositide or NMY-2 labeled (Fig. 8ABC). Moreover, The second and third cell divisions establishing the dorsal-ventral axis are also reconstructed virtually by the embryo with NMY-2 labeled (Fig. 8C), similar to the phosphoinositide-labeled embryos (Fig. 6AB, Movie S6). This suggests NMY-2 as an alternative fluorescently-labeled molecule for time-lapse cell membrane segmentation, accompanied by quantifiable spatiotemporal dynamics on the cell membrane.

**Fig. 8.**
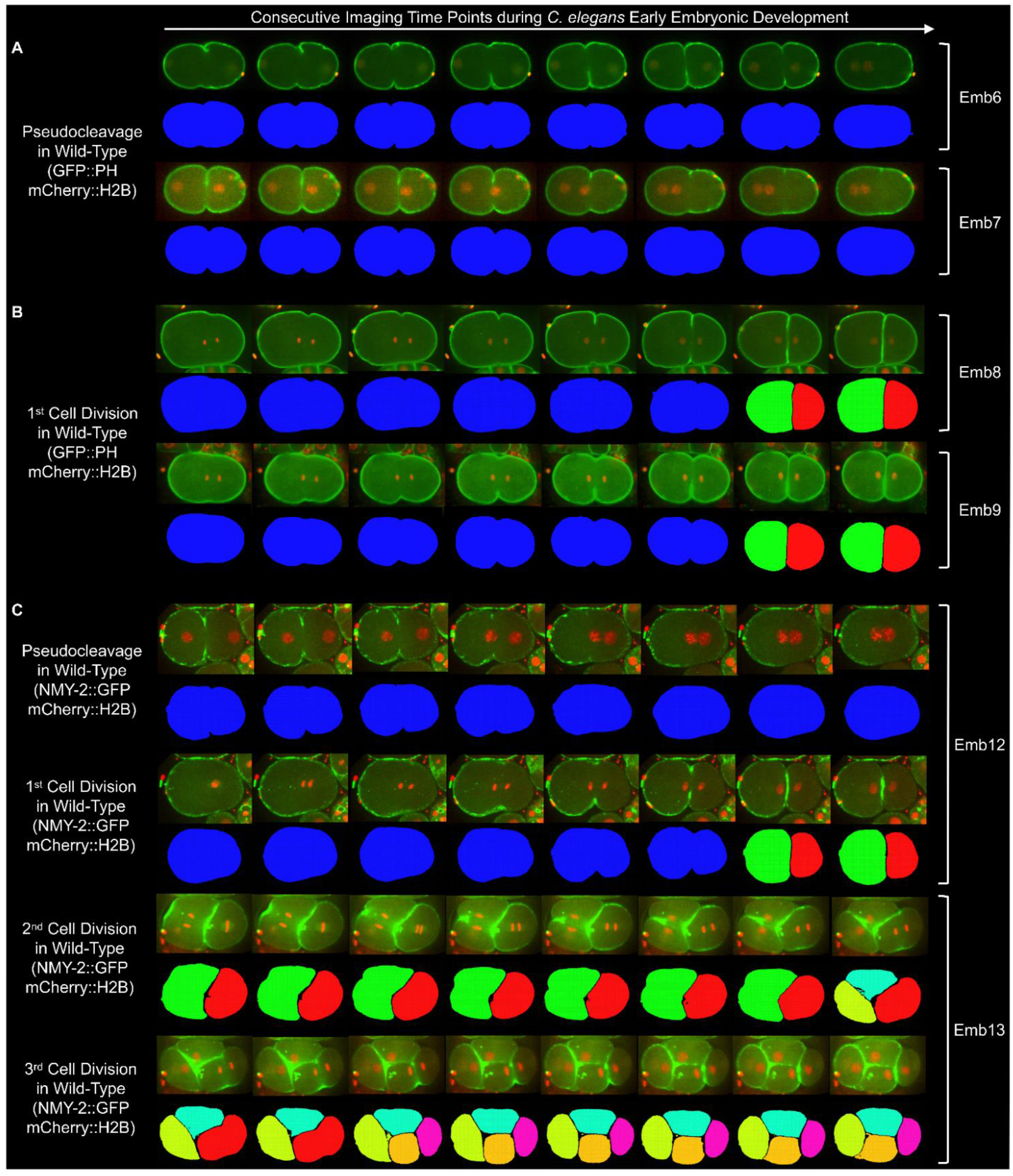
The segmentation results for the wild-type embryos with alternative fluorescently-labeled molecules (green fluorescence). (**A**) Cell shape reconstruction for the postfertilization pseudocleavage with the fluorescence labeling phosphoinositide on the cell membrane (GFP::PH). (**B**) Cell shape reconstruction for the first cell division with the fluorescence labeling phosphoinositide on the cell membrane (GFP::PH). (**C**) Cell shape reconstruction for the postfertilization pseudocleavage and first to third cell divisions with the fluorescence labeling NMY-2 on the cell membrane (NMY-2::GFP). For (**A**)(**B**)(**C**), cell nuclei are fluorescently labeled (red fluorescence) by mCherry-tagged histone (mCherry::H2B).

We further tested if segmentation is also applicable to embryos with genetic backgrounds different from the wild-type one, so as to help phenotype identification and mechanism discovery. To this end, we generated *C. elegans* embryos with knockdown (RNAi) of three well-known cytokinesis-associated genes: *nop-1* (mediating cortical recruitment of Rho effectors and promoting Rho activation), *spd-1* (participating in mitotic spindle midzone assembly), and *par-3* (controlling cell polarity and asymmetric partitioning of cytoplasm) (Tse *et al*. 2012, Verbrugghe *et al*. 2004, Cuenca *et al*. 2003). Remarkably, segmentation performs well with clear cell membrane dynamics digitalized. Although all embryos complete the first cell division successfully, a spectrum of morphological defects appears during the postfertilization pseudocleavage before it. In both the *nop-1*^-^ and *spd-1*^-^ embryos, the pseudocleavage proceeds with the locally folded cell membrane unlike the globally smooth one in the wild-type embryo (Fig. 8A, Fig. 9AB); in the *par-3*^-^ embryo, the cell membrane folding is even more severe, forming bubble-like structures transiently (Fig. 9C). The successful segmentation paves the way for high-throughput gene function discovery by artificial perturbation, cell membrane morphology reconstruction, and quantitative phenotype identification in the future.

**Fig. 9.**
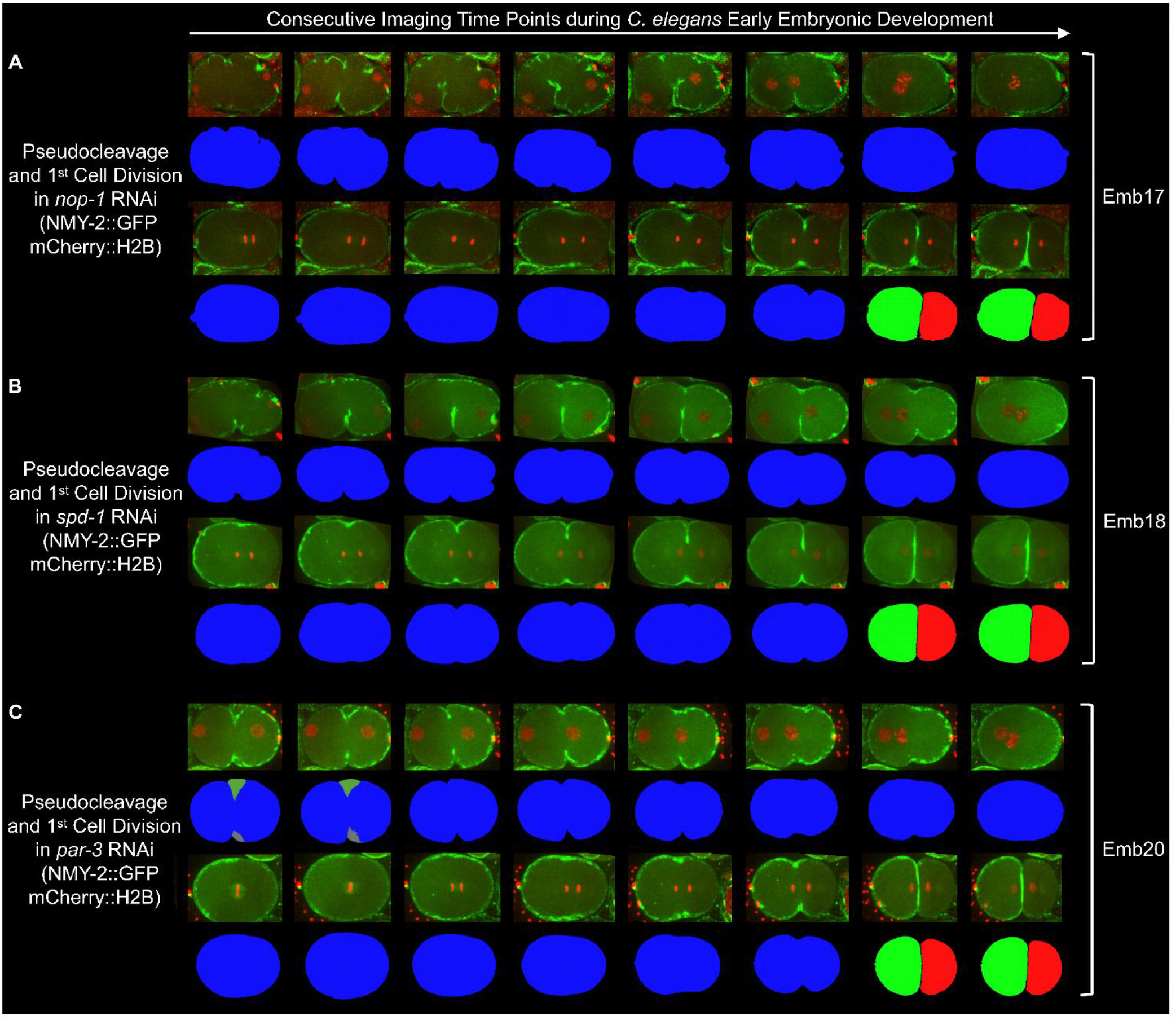
The segmentation results for the postfertilization pseudocleavage and first cell division in RNAi-treated embryos with fluorescently-labeled NMY-2 (green fluorescence). (**A**) The embryo with *nop-1* knockdown. (**B**) The embryo with *spd-1* knockdown. (**C**) The embryo with *par-3* knockdown. For (**A**)(**B**)(**C**), cell nuclei are fluorescently labeled (red fluorescence) by mCherry-tagged histone (mCherry::H2B).

## 4. Conclusion

Quantitative and automatic reconstruction of time-lapse 3D cell shapes with fluorescently-labeled cell membranes is challenging, especially for a developing embryo, in which cells undergo rapid division and migration frequently coupled with cell fate specification and cell shape deformation. Such a challenge is even more severe when the fluorescence images exhibit a low SNR because of various experimental protocols (*e.g*., strains) and purposes (*e.g*., observing short-term or long-term biological processes) (Fig. S3). In this paper, we successfully segmented the time-lapse 3D images of five *C. elegans* wild-type embryo samples with all cell identities traced but with a relatively low SNR in their cell membrane fluorescence images, which had failed to be segmented by state-of-the-art algorithms. Such successful segmentation is realized by a newly-devised framework, *EmbSAM*, that contains a deep-learning-based denoising module and a watershed module followed by *SAM*, which allows accurate and robust reconstruction of the cell shapes from multiple developing embryos (Fig. 1, Fig. 2, Fig.3). With an exceptional temporal resolution as high as 10-second in two of the embryos, the results allow examination of the instantaneous change in cellular behaviors during rapid cell division of embryogenesis. For example, the detailed cytokinesis and fast directional cell motion during each cell division with known cell identities, cell lineages, and cell fates; upon those cell divisions, morphogenesis including body axes establishment and spatial reorganization for gastrulation is illustrated in 3D, along with quantitative cellular properties like cell division axis reorientation and cell surface area distribution presented in the time course (Fig. 4, Fig. 5, Fig. 6, Fig. 7). All the reconstructed time-lapse 3D cell shapes at 10-second intervals and calculated shape features (*incl*., cell volume, cell surface area, and cell-cell contact area) are publically available in the data format of software *ITK-SNAP-CVE* and website https://bcc.ee.cityu.edu.hk/CMOS/EmbSAM (Yushkevich *et al*. 2016, Guan *et al*. 2023).

Given the high segmentation accuracy and robustness of the *EmbSAM* framework, it could be applied not only to the wild-type embryos but also to the perturbed ones, such as the one curated with external compression, laser ablation, and RNA interference, to uncover how a developing embryo coordinates cellular behaviors (*e.g*., cell division and coupled motion) to enable the faithful formation of tissues or organs (Jelier *et al*. 2016, Guan *et al*. 2019, Kuang *et al*. 2022, Van Bavel *et al*. 2023, Pimpale *et al*. 2020, Hsu *et al*. 2023). With fifteen more *C. elegans* wild-type and RNAi-treated embryos imaged, cell membrane segmentation is proved to be feasible with fluorescence labeling on alternative molecules, including both the homogeneously-distributed one (phosphoinositide) and the heterogeneously-distributed one (NMY-2), where the latter one (profiles biologically significant dynamics, *i.e*., cell cortex fluidity associated with cell division and coupled motion) (Sugioka *et al*. 2018, Pimpale *et al*. 2020, Middelkoop *et al*. 2024) can replace the traditional marker for cell membrane labeling and leave more fluorescence channels for monitoring the dynamics of other molecules simultaneously; such monitoring can be customized for specific developmental stages from fertilization to late and even post embryogenesis, for the whole body or tissue/organ of interest (*e.g*., ACT-5 for monitoring the lumenal formation in intestinal cells) (Fig. S15) and at flexible intervals (*e.g*., down to 2-second) (Fig. S16) (Gobel et al. 2004, MacQueen *et al*. 2005). Following this paradigm, more molecule dynamics in time-lapse format can be collected quantitatively by strain crossing, fluorescence imaging, and cell segmentation in a high-throughput manner.

The fluorescence images of dozens of *C. elegans* embryos and the reconstructed cell shapes in 2D and 3D at high temporal resolutions in this work enable detailed analysis of cellular behaviors at the systematic level, which facilitates the construction of new theoretical models that help predict what’s going on in reality. For instance, the cell shape change during fast cell division and cytokinesis can help understand the cell membrane mechanics, providing a reference for testing various cell membrane models established previously (Ma *et al*. 2014, Kuang *et al*. 2022, Cuvelier *et al*. 2023, Ichbiah *et al*. 2023). Moreover, such embryo-wide cell shapes can be used to reversely infer the intracellular and intercellular mechanical properties over development, which are usually hard to measure directly (Xu *et al*. 2018, Guan *et al*. 2023, Yamamoto *et al*. 2023, Ichbiah *et al*. 2023). When replacing the fluorescent marker used for labeling cell membranes with those specific for other cellular or subcellular compartments (*e.g*., E-cadherin and F-actin), the dynamics of their related cellular behaviors, such as cell adhesion and cell stiffness, can be examined with an exceptional temporal resolution, *i.e*., at a 10-second or a shorter interval (Yamamoto *et al*. 2017, Fujita *et al*. 2012). Such real-time dynamics of cellular or subcellular behaviors not only help understand the biological regulation *in vivo*, but also facilitate the establishment of a reliable computational model that simulates developmental control *in silico* and permits virtual experiments for mechanism discovery (Hubatsch *et al*. 2019, Cuvelier *et al*. 2023, Kuang *et al*. 2023). In the future, the *EmbSAM* framework could be used to analyze datasets with fluorescently-labeled cell membranes beyond the ones analyzed in this work, such as those of ascidian, fruit fly, zebrafish, and mouse, so as to broaden the cell shape data in various biological context (Guignard *et al*. 2020, Stegmaier *et al*. 2016, Khan *et al*. 2014).

## Availability and implementation

The data is available at https://doi.org/10.6084/m9.figshare.26955949.v1.

The code is available at https://github.com/cunminzhao/EmbSAM.

The website is available at https://bcc.ee.cityu.edu.hk/cmos/embsam

## Acknowledgement

We would like to thank Dr. Yu Chung Tse and Dr. Xiangchuan Wang from SUSTech Core Research Facilities at Southern University of Science and Technology and Prof. Chuanfu Dong at Beijing Normal University for providing technical support on spinning disk confocal imaging as well as the comparison of fluorescence patterns between wild-type and RNAi-treated *C. elegans* embryos with various markers. We would also like to thank Dr. Dongying Xie, Yiming Ma, and Gefei Huang at Hong Kong Baptist University for the fruitful discussion.

