## Supplement for "*EmbSAM*: Cell boundary localization and Segment Anything Model for fast images of developing embryos"

### Supplemental Figures

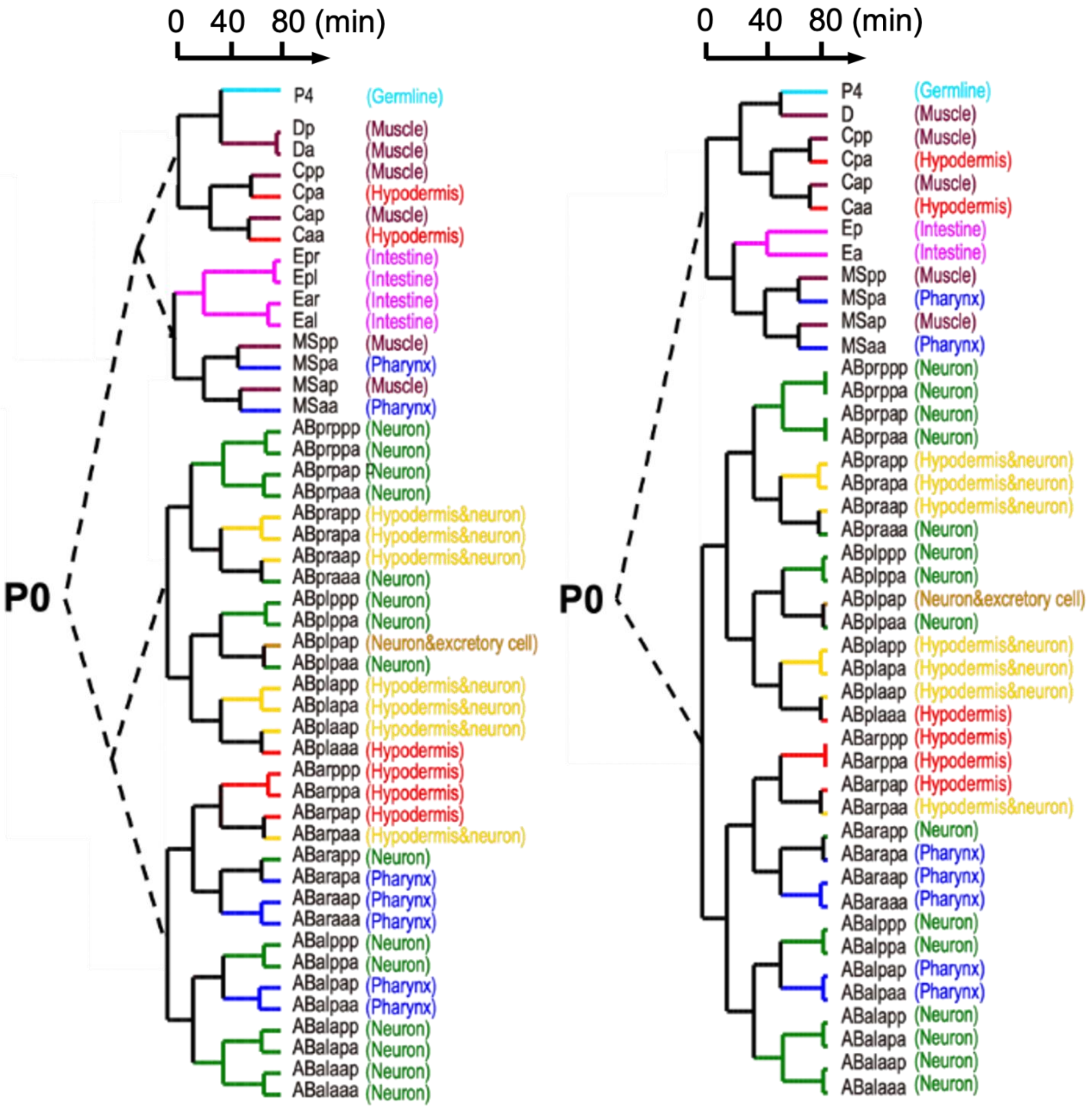


**Fig. S1.** The cell lineage tree of *C. elegans* early embryogenesis (exemplified by the embryo samples "Emb1" and "Emb2" from left to right), with the unique cell identities and primary cell fates labeled (Sulston *et al.* 1983, Ho *et al.* 2015).

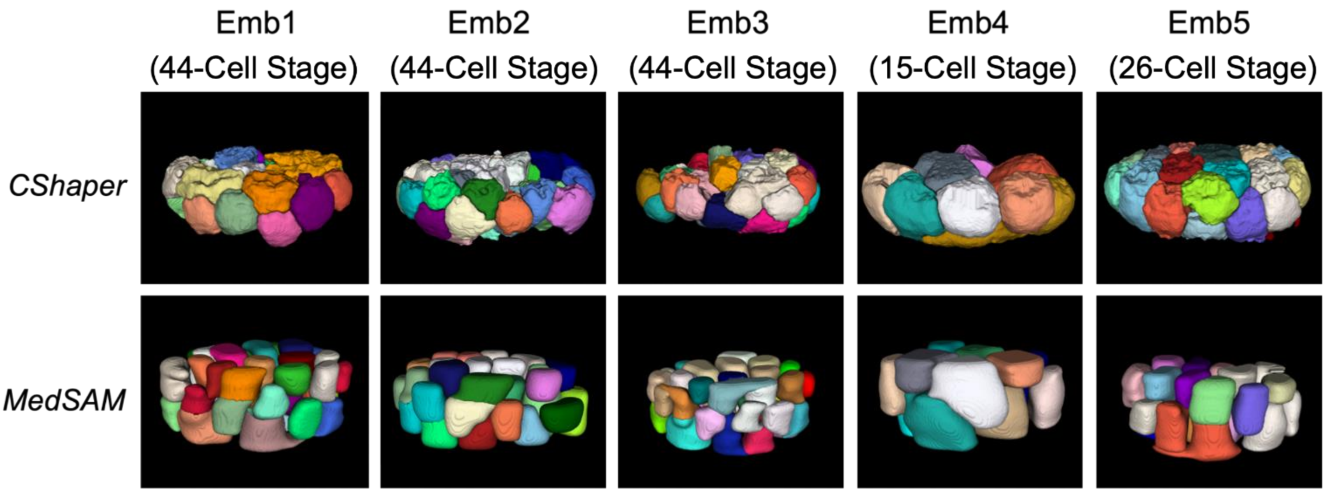


**Fig. S2.** The poor segmentation results (exemplified by the embryo samples "Emb1" to "Emb5") of *CShaper* and *MedSAM* at different developmental stages (Cao *et al.* 2020, Ma *et al.* 2024), revealed by the coarse and uncompacted cell shapes. Shown are side views with the anterior of the embryo to the right.

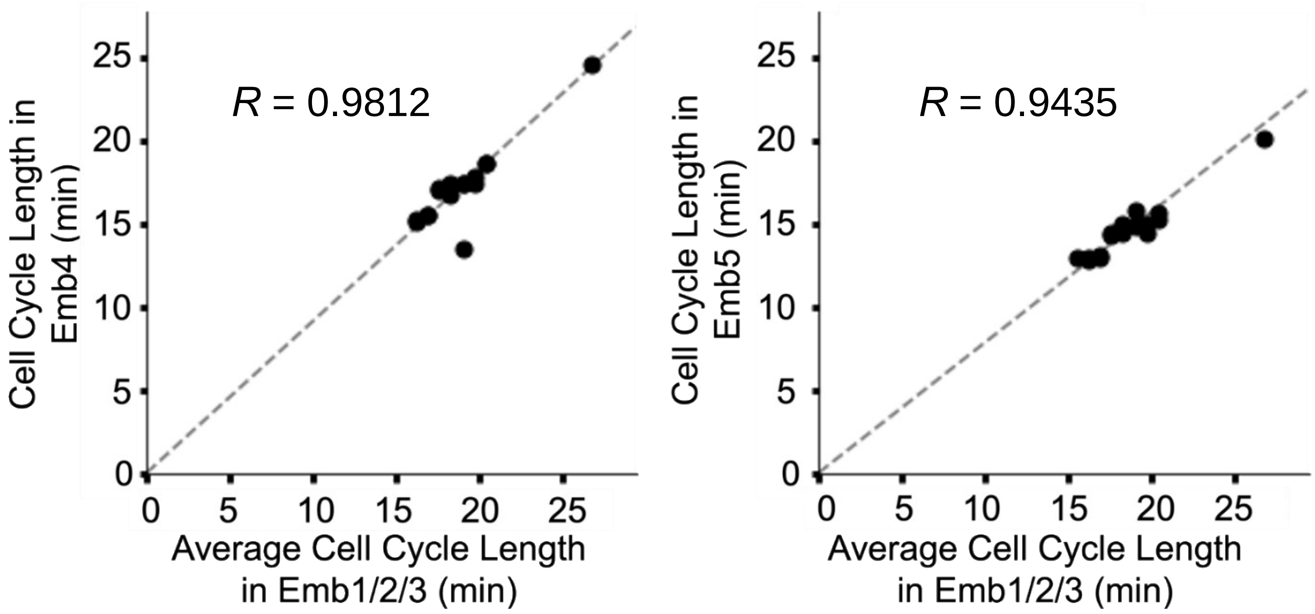


**Fig. S3.** The proportional relationship between the cell cycle lengths in the embryo samples "Emb1" to "Emb3" and the embryo samples "Emb4" (left) and "Emb5" (right), with the goodness of fit labeled in the top left corners.

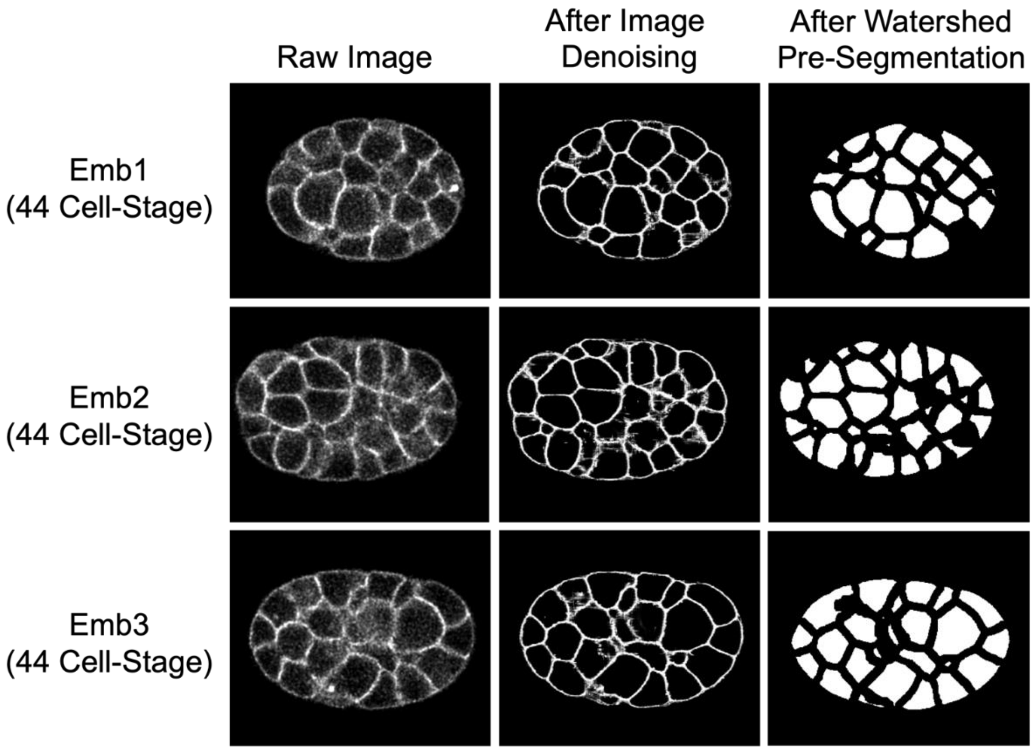


**Fig. S4.** The raw and processed images (exemplified by the embryo samples "Emb1" to "Emb3") of *EmbSAM* before the *SAM* segmentation.

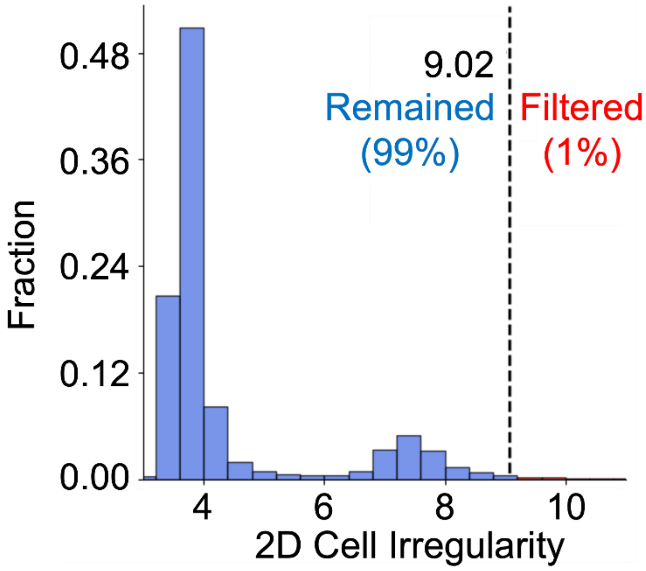


**Fig. S5.** The distribution of 2D cell irregularity generated from the manually-annotated cell regions in previous works (Table S1) (Cao *et al.* 2020, Guan *et al.* 2023), with the minimum of the 1% largest values set as the threshold for filtering abnormal 2D cell regions.

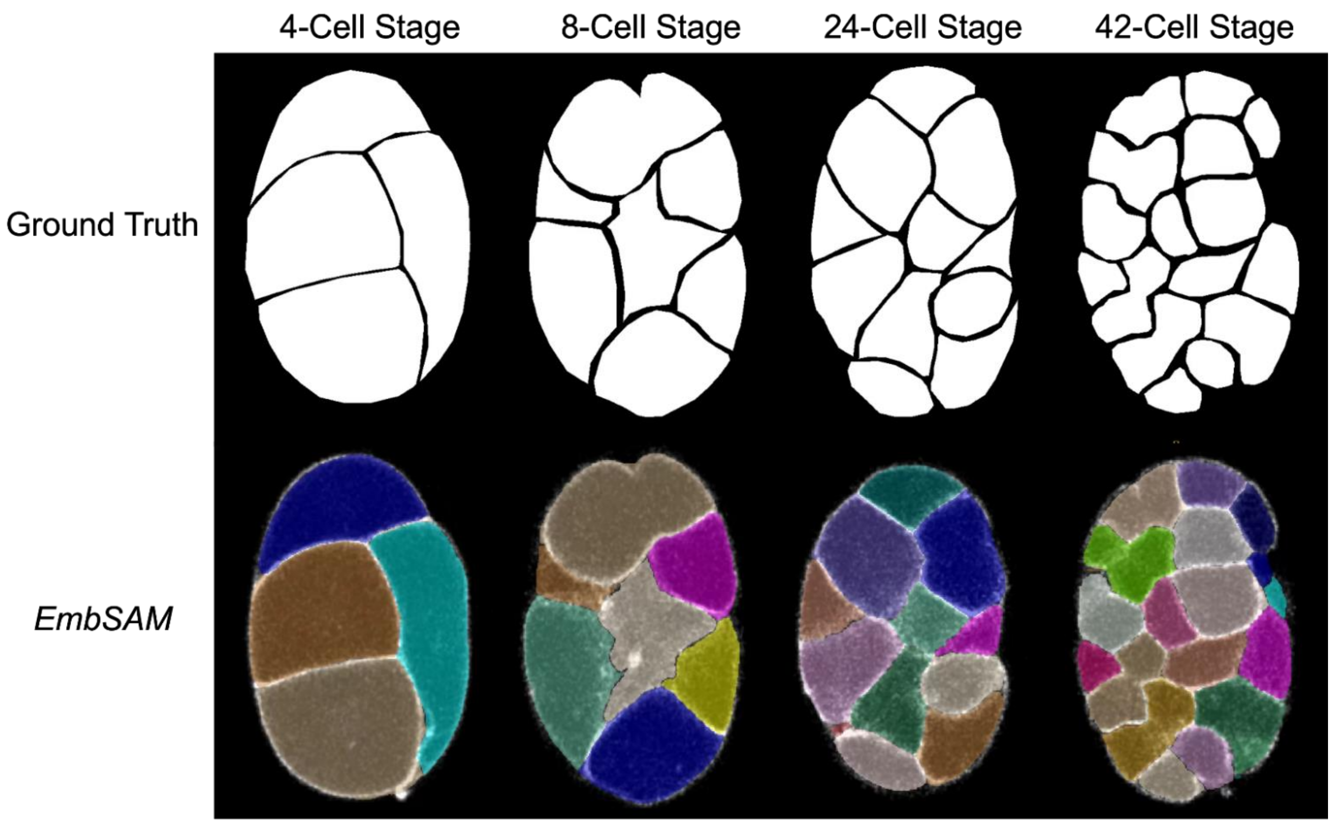


**Fig. S6.** The segmentation results (exemplified by the embryo sample “Emb3”) of *EmbSAM* near the middle focal plane and at different developmental stages.

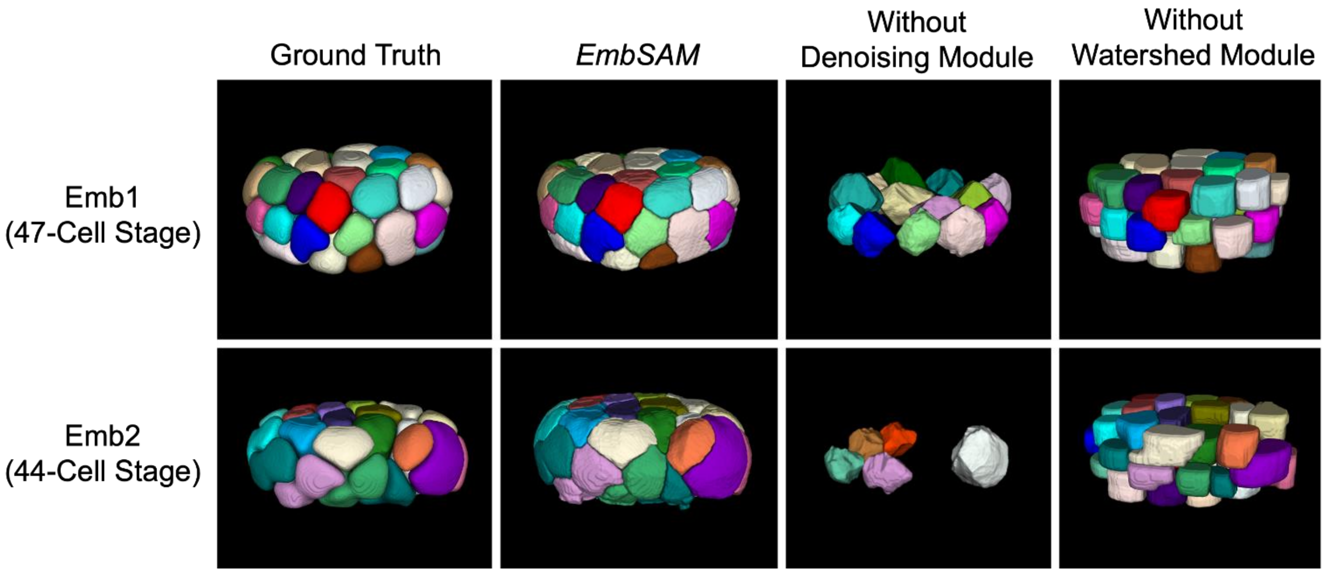


**Fig. S7.** The ground truth and segmentation results (exemplified by the embryo samples “Emb1” and “Emb2”) of *EmbSAM* and its truncated versions with the denoising module and watershed module removed.

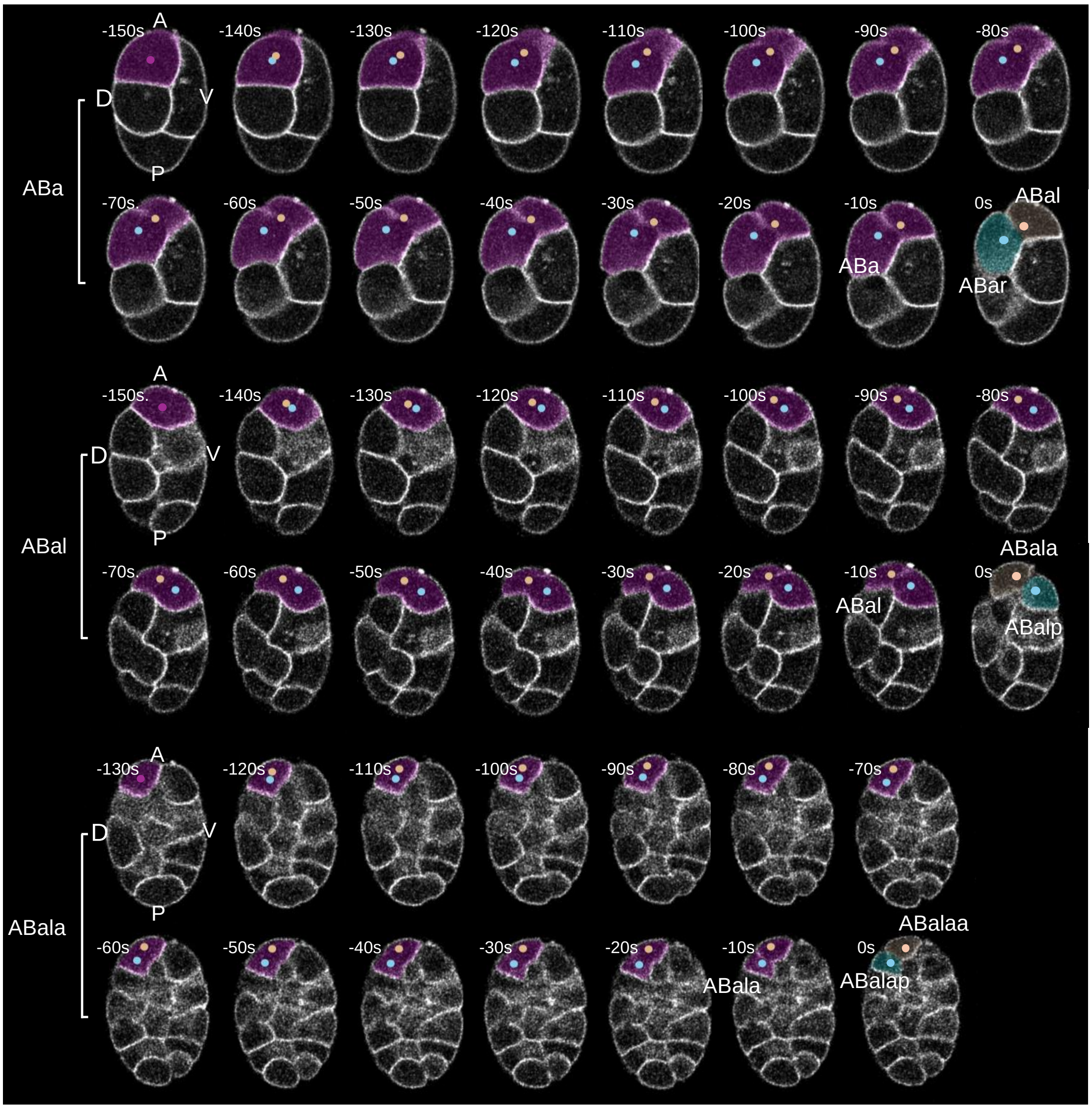


**Fig. S8.** The segmentation results (exemplified by the embryo sample "Emb5") of *EmbSAM* for the ABa, ABal, and ABala cell divisions (from top to bottom) at 10-second intervals, viewed in the shooting direction and highlighted by the dotted cell nuclei and masked cell membranes. Shown are lateral views with the anterior of the embryo to the top. In each subfigure, the absolute developmental time is shown, with the moment of complete cell membrane separation as time zero.

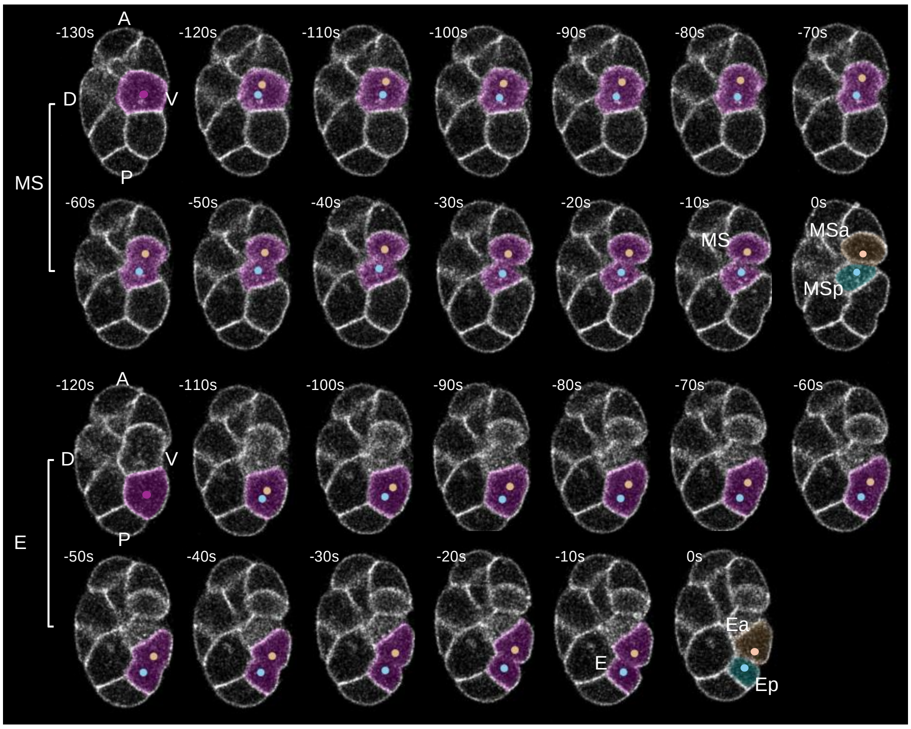


**Fig. S9.** The segmentation results (exemplified by the embryo sample "Emb5") of *EmbSAM* for the MS and E cell divisions (from top to bottom) at 10-second intervals, viewed in the shooting direction and highlighted by the dotted cell nuclei and masked cell membranes. Shown are lateral views with the anterior of the embryo to the top. In each subfigure, the absolute developmental time is shown, with the moment of complete cell membrane separation as time zero.

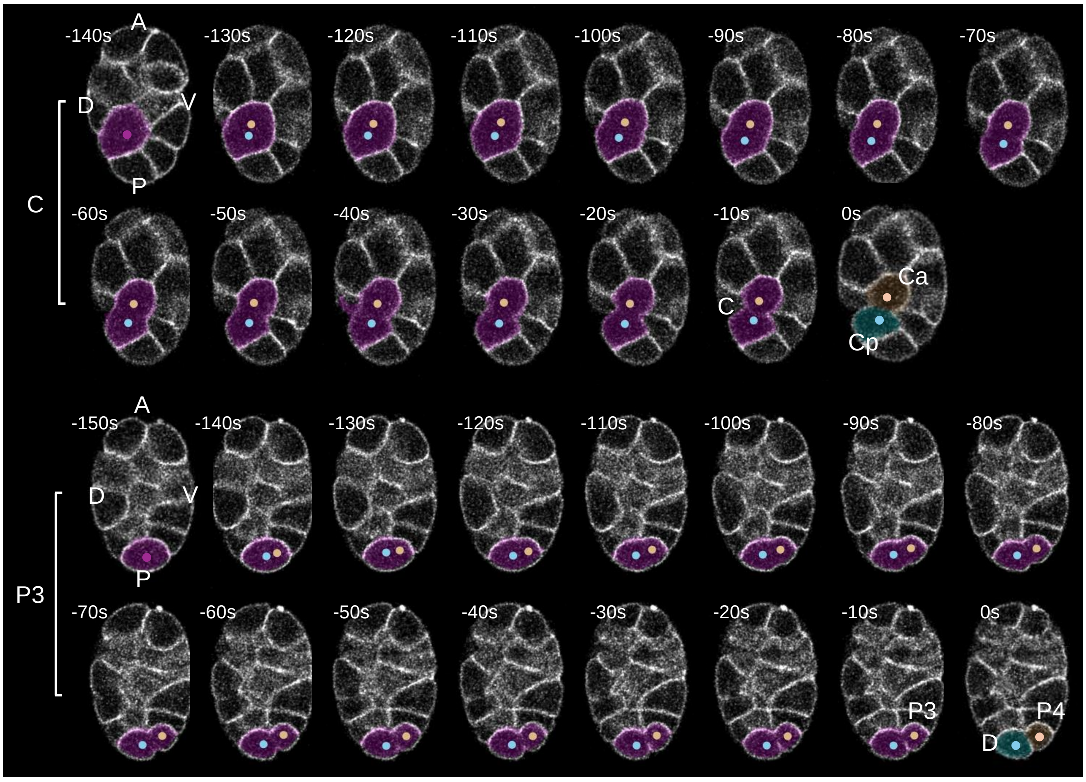


**Fig. S10.** The segmentation results (exemplified by the embryo sample "Emb5") of *EmbSAM* for the C and P3 cell divisions (from top to bottom) at 10-second intervals, viewed in the shooting direction and highlighted by the dotted cell nuclei and masked cell membranes. Shown are lateral views with the anterior of the embryo to the top. In each subfigure, the absolute developmental time is shown, with the moment of complete cell membrane separation as time zero.

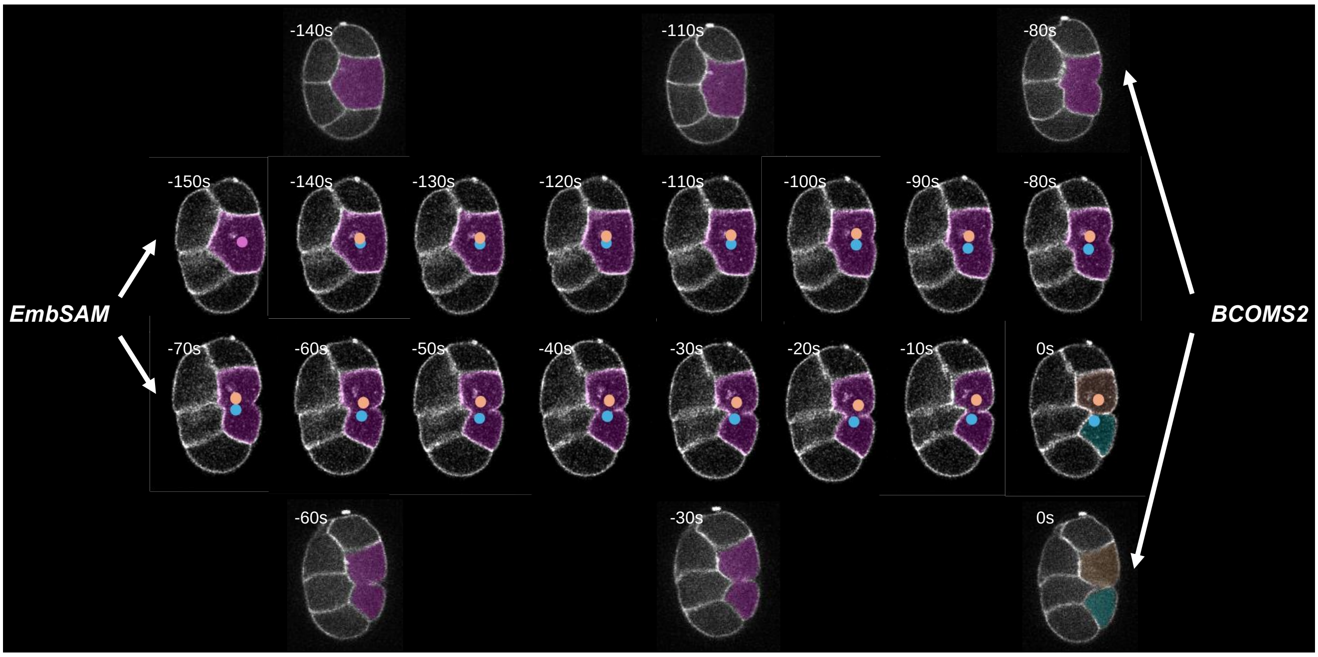


**Fig. S11.** The segmentation results (exemplified by the embryo sample “Emb5”) of *EmbSAM* and *BCOMS2* (Azuma *et al.* 2023) for the EMS cell division at 10-second intervals, viewed in the shooting direction and highlighted by the dotted cell nuclei and masked cell membranes. Shown are lateral views with the anterior of the embryo to the top. In each subfigure, the absolute developmental time is shown, with the moment of complete cell membrane separation as time zero.

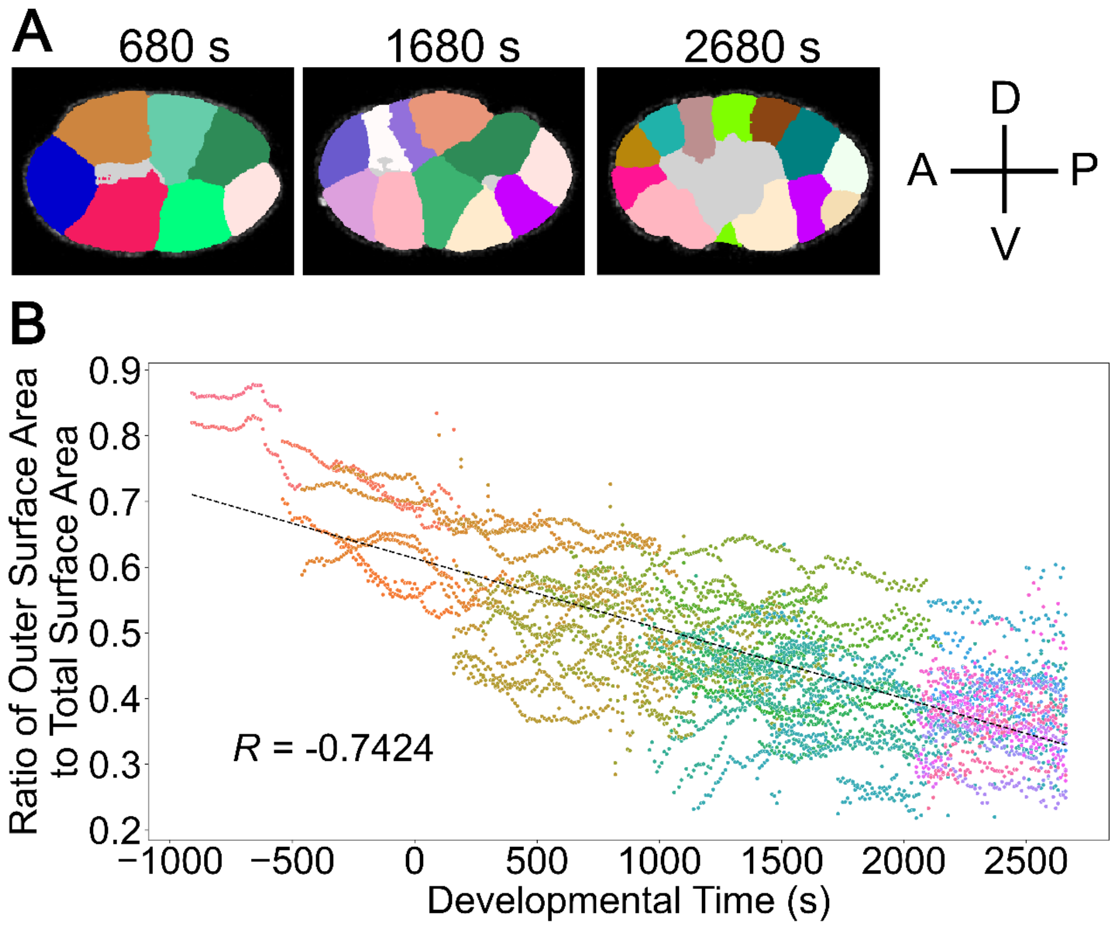


**Fig. S12.** The segmentation results (exemplified by the embryo sample “Emb5”) of *EmbSAM* for the spatial reorganization for gastrulation. (**A**) Crosssection viewed in the shooting direction, with the cells regularly attached to the eggshell colored differentially and the region inside colored in gray. (**B**) Linear relationship between the ratio of outer surface area (considering all cell regions in the embryo samples “Emb4” and “Emb5”) and developmental time, with the linear correlation coefficient labeled in the bottom left corner. For (**A**)(**B**), the absolute developmental time is shown, with the last moment of the 4-cell stage as time zero.

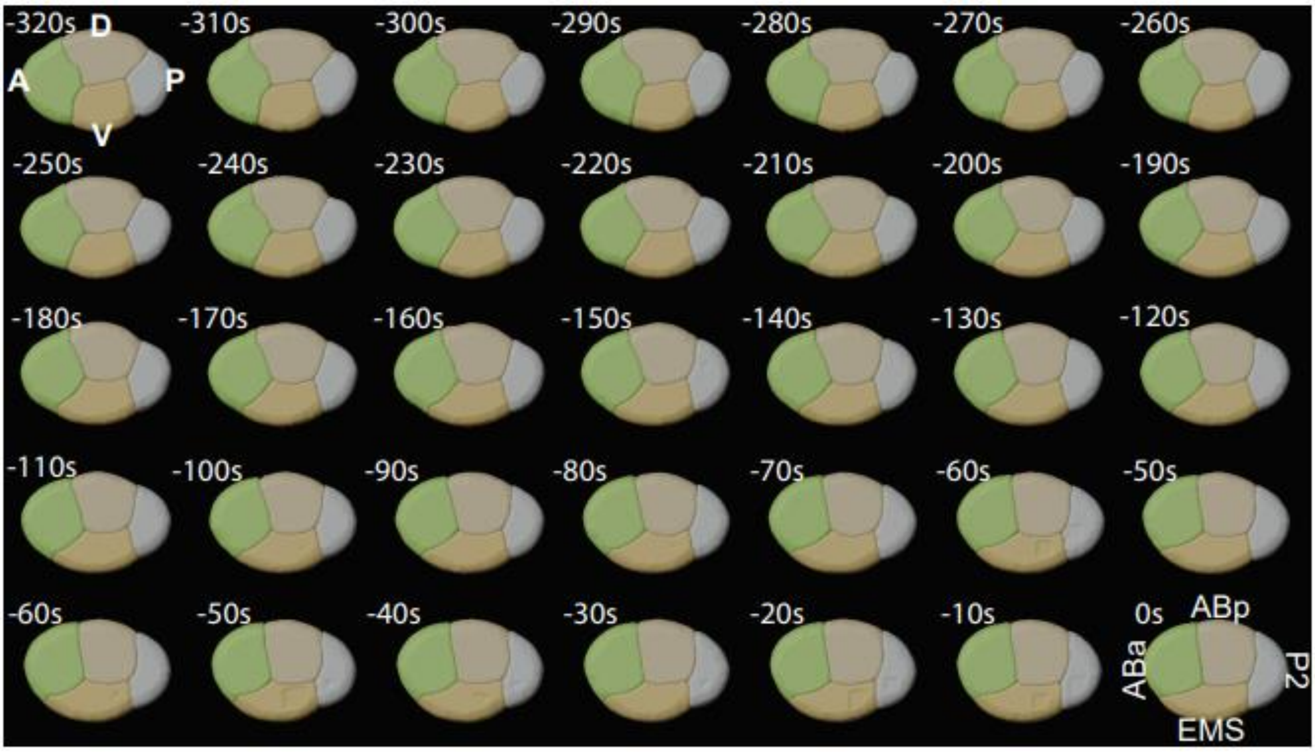


**Fig. 13.** The segmentation results (exemplified by the embryo sample "Emb4") of *EmbSAM* for the late 4-cell stage at 10-second intervals, viewed in the shooting direction. Shown are lateral views with the anterior of the embryo to the left. In each subfigure, the absolute developmental time is shown, with the last moment of the 4-cell stage as time zero.

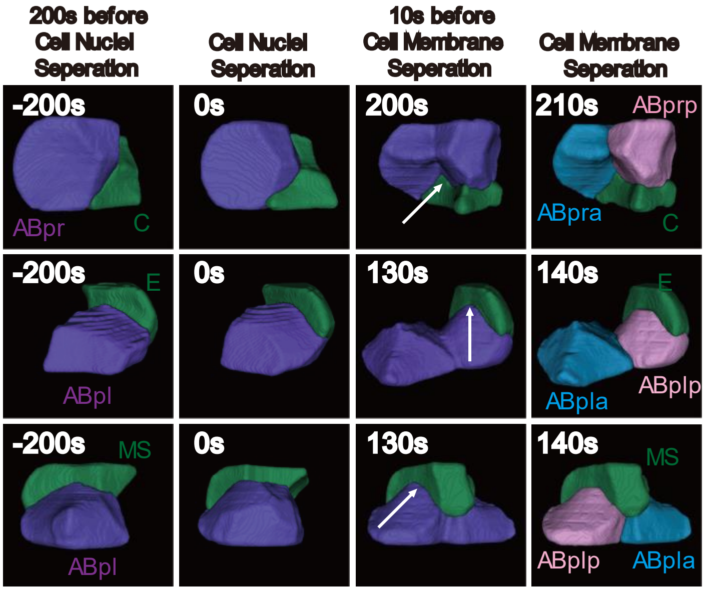


**Fig. 14.** The segmentation results of *EmbSAM* for the drastic shape dynamics of dividing cells (purple), their neighbors (green), and newborn daughters (pink and blue) at 10-second intervals, illustrated by snapshots for three cell division events (from top to bottom). In each subfigure, the absolute developmental time is shown, with the moment of complete cell nuclei separation as time zero.

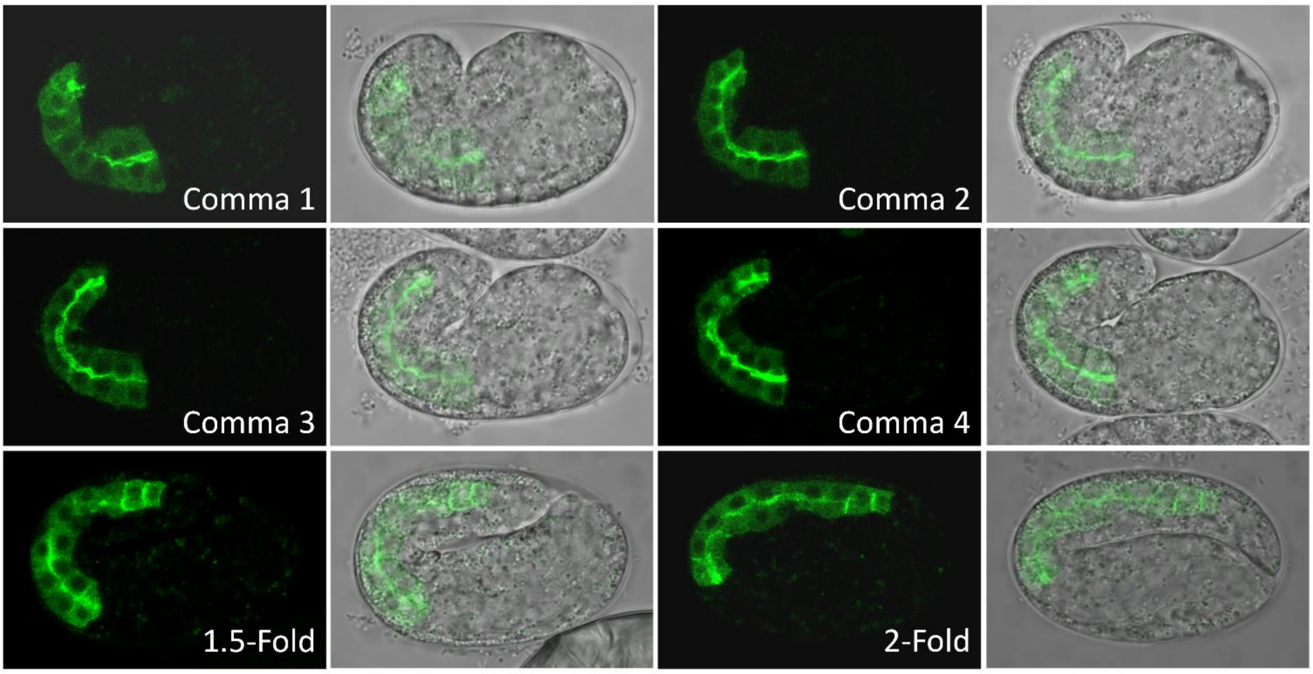


**Fig. S15.** The single-layer fluorescence (left; globally enhanced) and bright-field (right; superimposed with original fluorescence) images for the wild-type embryos at the comma (intermediate stages indicated by 1 to 4), 1.5-fold, and 2-fold stages. Intestinal cells and their lumen are fluorescently labeled (green fluorescence) by ACT-5::GFP, which tags the microvillar actin bundle and has substantially high accumulation in the apical surface of intestinal cells.

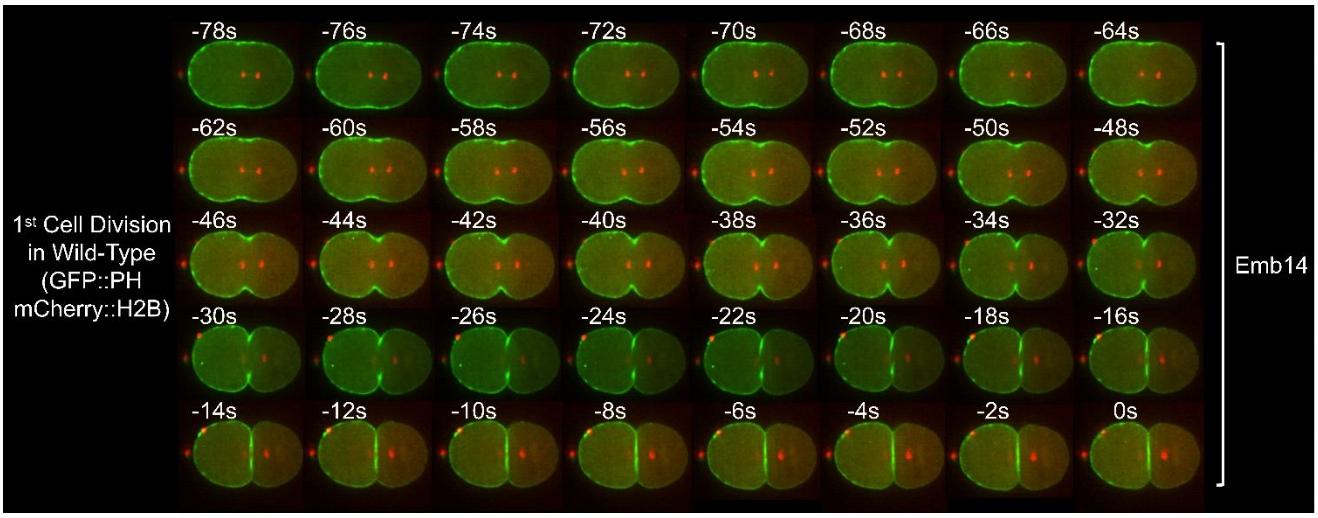


**Fig. S16.** The projected fluorescence images for the wild-type embryo during the first cell division. In each subfigure, the absolute developmental time is shown, with an arbitrarily selected time point when the cell division is surely completed as time zero. Cell membrane is fluorescently labeled (green fluorescence) by GFP-tagged phosphoinositide (GFP::PH). Cell nuclei are fluorescently labeled (red fluorescence) by mCherry-tagged histone (mCherry::H2B).

### Supplemental Tables

**Table S1.** The number of manually-annotated cell regions in previous works (Cao *et al.* 2020, Guan *et al.* 2023).

**Table S2.** The information on embryos with alternative fluorescently-labeled molecules and different genetic backgrounds.

### Supplemental Movies

All time-lapse 3D movies are generated from segmented fast-imaging embryos at 10-second intervals. The 3D images in the movies are rendered with the software Fiji and Blender (Schindelin *et al.* 2012, Blender Online Community 2024).

**Movie S1.** The time-lapse 3D segmentation results of the embryo sample "Emb4", viewed along the left-right and dorsal-ventral axes.

**Movie S2.** The time-lapse 3D segmentation results of the embryo sample "Emb5", viewed along the left-right and dorsal-ventral axes.

**Movie S3.** The user instruction of the *ITK-SNAP-CVE* software for embryos digitized at 10-second intervals.

**Movie S4.** The user instruction of the *CMOS* website for embryos digitized at 10-second intervals.

**Movie S5.** The time-lapse 3D segmentation results of the EMS cell division (exemplified by the embryo sample "Emb5") from the 4- to 8-cell stages, viewed along the left-right axis.

**Movie S6.** Time-lapse 3D segmentation results of the 2- to 4-cell stages with the establishment of the dorsal-ventral axis and lifespan-dependent cell shape changes (exemplified by the embryo sample "Emb4"), viewed along the left-right and dorsal-ventral axes.

**Movie S7.** Time-lapse 3D segmentation results of the 6-cell stage with the establishment of the left-right axis and skewed ABa/ABp cell division orientations (exemplified by the embryo sample "Emb4"), viewed along the left-right and dorsal-ventral axes.

**Movie S8.** Time-lapse 3D segmentation results of the 7-cell stage with the long-range ABpl cell migration (exemplified by the embryo sample "Emb4"), viewed along the left-right and dorsal-ventral axes.
